# The alphavirus TF protein plays a critical role in promoting viral propagation by altering cell-cell boundaries

**DOI:** 10.1101/2023.07.24.550336

**Authors:** Pushkar Tatiya, Ramesh Kumar, Debajit Dey, Yashika Ratra, Syed Yusuf Mian, Subhomoi Borkotoky, Shaileshanand Jha, Bhawna, Sanchi Arora, Kirti Suhag, Soumen Basak, Manidipa Banerjee

## Abstract

During cellular infections, alphaviruses produce a 6 kDa membrane protein (6K) and a transframe variant (TF) of 6K through ribosomal frameshifting. The role of these proteins in alphavirus biology is largely unknown. Here, we show that TF of Chikungunya virus plays a unique and critical role in promoting virus propagation by manipulating host cell boundaries. TF achieves this by targeting and downregulating human Scribble, a component of the polarity complex that controls cellular morphology and apoptosis, leading to its punctate localization, ubiquitination, and subsequent degradation via the proteasomal pathway. The TF-Scribble interaction is mediated by a newly discovered PDZ-domain binding motif (PBM) in TF, with the PDZ domains of Scribble, as demonstrated by co-localization during viral infection and further by co-immunoprecipitation and proteomics analysis. This PBM is unique to TF and is absent in 6K; thus, 6K cannot modulate Scribble localization in infected cells. shRNA-mediated knockdown of Scribble potentiates CHIKV propagation highlighting the requirement of TF for the downregulation of Scribble. While wildtype CHIKV drastically alters cell-cell boundaries, a mutated version of CHIKV generated through reverse genetics (ΔTF CHIKV), which produces 6K but not TF, is unable to trigger similar morphological alterations due to its inability to engage Scribble, ultimately resulting in suboptimal virus propagation. A cryoEM reconstruction of ΔTF CHIKV at a global resolution of 3.6 Å, one of the highest reported among all alphavirus structures, indicates that the assembly of glycoprotein spikes and nucleocapsid is unaffected. Thus, the loss of TF affects the ultimate stage of cellular egress in the virus life cycle, and not earlier stages such as entry, replication or assembly. Our work thus establishes a novel function for the TF component of alphaviruses in modulating host morphology, potentiating virus transmission, and also distinguishes the functionality of TF from that of 6K in alphavirus biology.

**Teaser:** A novel functionality of Chikungunya Virus in modulating cell boundaries to allow efficient virus propagation

## Introduction

Chikungunya Virus (CHIKV), a member of the alphavirus genus in the *Togaviridae* family, is a mosquito-borne human pathogen that affects almost 100 countries and causes ∼3,000,000 infections per year globally (1). There are no targeted therapeutics or licensed preventive strategies for CHIKV, and the treatment is primarily symptomatic (2, 3). While the symptoms caused by this virus are self-limiting, there can be severe complications such as chronic, arthritis-like symptoms; and infections in the very young and elderly populations can be fatal. A clearer understanding of the functions of CHIKV-encoded proteins and the identification of their host interaction partners are crucial for the development of targeted therapeutics or attenuated vaccine candidates (4).

The CHIKV capsid, like that of all alphaviruses, is a 60-70 nm particle with T=4 icosahedral geometry (5–7). The outer layer is composed of 240 heterodimers of envelope glycoproteins E1 and E2, embedded in a host-derived lipid envelope. While E2 is responsible for association with the host cellular receptor(s), E1 orchestrates the fusion of the viral and host lipid membranes (4). The inner layer of the particle consists of 240 copies of a multifunctional capsid protein (C) which encapsulates the genome (8). Another envelope protein, E3, promotes the association between E1 and E2 during virus assembly (9).

The functionalities of the major structural proteins of CHIKV – E1, E2, E3, and C - are well investigated and established. However, a fifth protein – the small peptide “6K”- remains relatively underexplored, and its role in the virus life cycle is unclear (4). It has been shown that recombinantly generated alphavirus 6K is an ion channel-forming protein (10, 11). 6K from Ross River Virus (RRV) and Barmah Forest Virus (BFV) form cation-specific channels in bilayer membranes in electrophysiology experiments (11). The expression of 6K in bacterial cells is detrimental to cell growth, consistent with 6K possessing membrane-disrupting activity (11, 12). Bacterially produced CHIKV 6K, in conjunction with the expression partner Glutathione-S-transferase (GST), forms heterogeneous pores in the membrane *in vitro* (10). Experiments in cellular overexpression models and *in vitro* liposome disruption studies have shown that CHIKV 6K tends to have a strong affinity for Endoplasmic Reticulum (ER) membranes and congregates in the ER. CHIKV 6K, as well as Sindbis Virus (SV) 6K, co-localize with the glycoprotein E2. SV 6K co-localizes with E2 in cytopathic vesicles on the way to the plasma membrane, which is the budding site for alphaviruses (10,13,14). It has been conjectured that 6K is required to facilitate concurrent transport of viral protein and genomic complexes and to promote association of glycoproteins during assembly (14), although there is no clear connection between this functionality and the ion-channel activity of 6K at present.

The role of 6K in the alphavirus life cycle has been further complicated by the discovery of a transframe variant “TF”, which is generated due to ribosomal slippage within the 6K gene. TF is produced at a 10-18 % frequency during viral protein translation (15, 16) and shares the majority of the primary amino acid sequence with 6K but varies in the C-terminal region. The production of TF results in premature termination of structural polyprotein translation; thus, E1 is not generated along with TF (Fig. 1A). Post-translational modification, specifically palmitoylation, of the Sindbis Virus (SV) TF, is necessary for its association with the plasma membrane (17). Alphavirus capsids may contain TF as a structural component, albeit in sub-stoichiometric quantities (15,18). Recent work has shown that, while recombinantly generated TF can also form ion channels in membranes, the unique C-terminal tail influences the overall conformation, resulting in a more open and less complex structure compared to 6K (19). The exact functionality of TF during alphavirus infections is not established; and it remains a mysterious entity like 6K. The functionalities attributed to 6K in many previous works may have been, to some limited extent, influenced by the coexistence of hybrid peptides containing partial sections of TF, due to the non-removal of the ribosomal slippage site within the 6K genomic sequence (10,17).

**Fig. 1.**
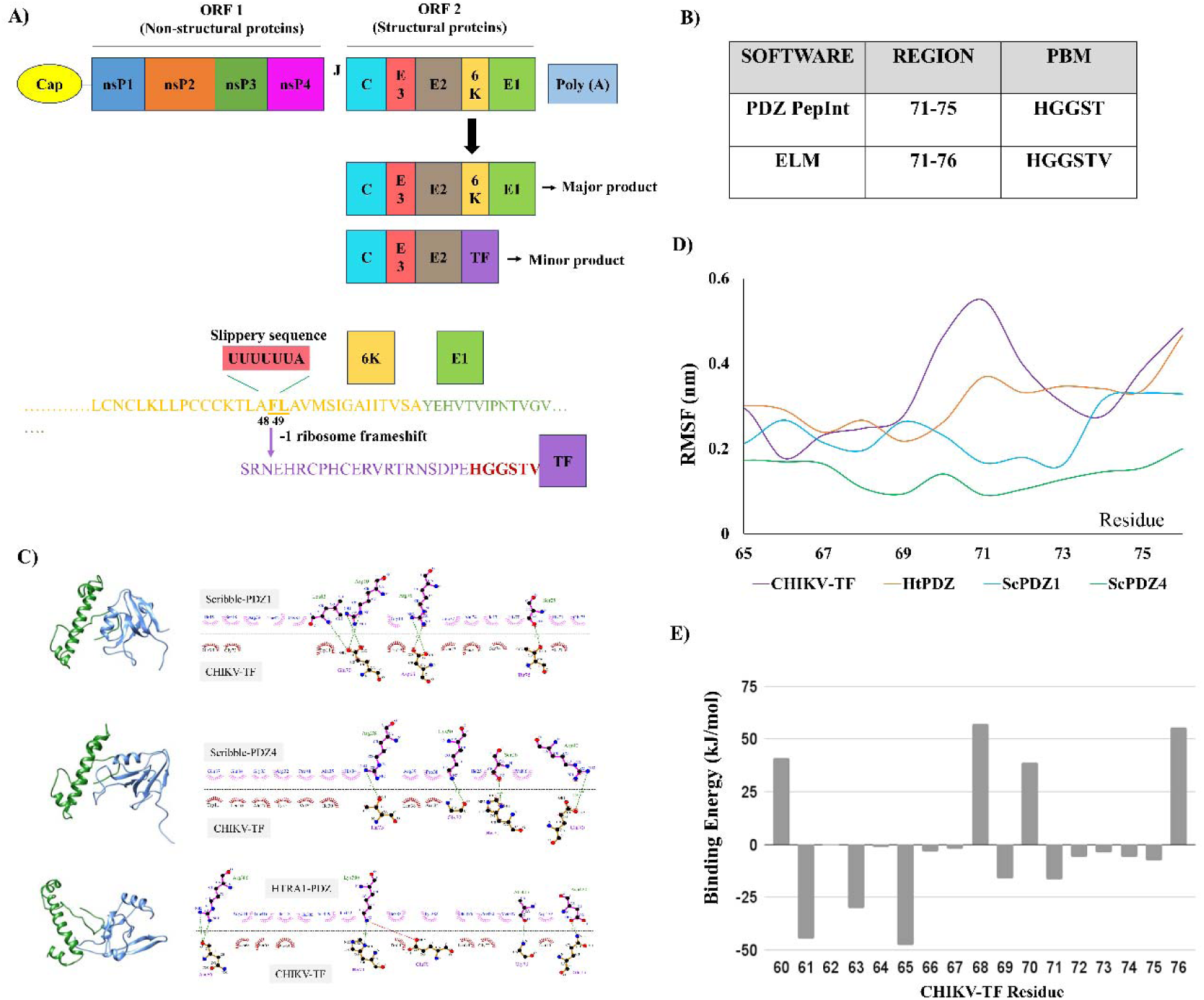
CHIKV TF as a binding partner for human Scribble. The CHIKV RNA genome has two open reading frames (ORFs) separated by a junctional region (J). The 5’ ORF (ORF 1) codes for four non-structural proteins (nsP1-4). The 3’ ORF (ORF 2) codes for 5 structural proteins, namely capsid (C), E3, E2, 6K, and E1. The 6K gene has a ribosomal slippery site which leads to the production of a transframe (TF) protein (Firth *et al.,* 2008) (upper panel, A). Schematic of 6K and TF sequences and the ribosome slip site (F at 48^th^ and L at 49^th^ position) (lower panel, A). Potential PDZ binding motifs (PBMs) present in TF as predicted by different softwares (B). Predicted interactions between TF and Scribble PDZ1, Scribble PDZ4, and HTRA1 (C). RMSF plot for TF interactions with Scribble and HTRA1 (D). Binding of TF C-terminal residues with ScPDZ1(E).

Upon a close examination of the CHIKV TF amino acid sequence, we noted that the unique region corresponding to the C-terminal region of TF contains a classical PDZ-domain binding motif (PBM). The acronym PDZ originates from the first three proteins discovered to contain this domain – the postsynaptic density protein 95 (PSD95), Drosophila disc large tumour suppressor (Dlg1), and zonula occludens-1 protein (ZO-1) (20, 21). PDZ domains are protein-protein interaction domains found in practically all life forms. *In silico* docking and simulation studies with the PBM at the C-terminus of TF suggested that a potential binding partner for this motif is Scribble, part of a conserved set of proteins that controls apico-basal as well as planar cell polarities and is also involved in the modulation of tissue morphogenesis and apoptosis (22,23). Although the involvement of Scribble in cellular infections by CHIKV or other alphaviruses is, to date, unknown, non-structural proteins of different viruses, such as Influenza Virus, Human T lymphotropic Virus-1 (HTLV-1), and Hepatitis C Virus (24), target scribble. In this study, we report that CHIKV infection results in more than 90% abrogation of Scribble protein. Scribble is ubiquitinated and marked for degradation within 24 hours of CHIKV infection in HEK293T cells *in vitro*. We present evidence for the involvement of TF in Scribble degradation, as transfection of TF or its production during CHIKV infection results in the sequestration of Scribble into discrete cytosolic puncta. TF also colocalizes in these puncta and can be immunoprecipitated with Scribble during CHIKV infection. The co-localization as well as pulldown of TF with Scribble can be abrogated by mutations within the PBM of TF, showing that the interaction between TF and Scribble is mediated by a PDZ-PBM association. The knockdown of Scribble results in a significant enhancement in CHIKV infectivity in cells. TF-mediated sequestration of Scribble accelerates virus propagation through drastic modification of the cellular boundaries. A version of CHIKV generated through reverse genetics, that does not produce TF, has reduced ability to engage with Scribble and cannot alter cellular boundaries. A cryoEM map of this mutated version of CHIKV clearly indicates that this particle is correctly assembled and therefore competent in all other ways to infect host cells but impeded in its propagation due to the lack of TF. Our work thus establishes the enigmatic TF protein as an essential factor for strategic manipulation of host cell morphology during alphavirus infections leading to enhanced pathogen survival and replication efficiency.

## Results

### The unique C-terminus of CHIKV TF contains a PDZ domain binding motif with potential for association with Scribble

The first 49 amino acids in the protein sequence of CHIKV 6K and TF are identical. This region is highly hydrophobic and capable of interacting with lipid membranes, as shown previously (10,15). A ribosomal -1 frameshift at a slippery sequence corresponding to residues 48-49 within the 6K gene results in the production of TF with a unique C-terminal region of 27 amino acid residues which is primarily hydrophilic (Fig. 1A). A motif search of this region indicated the presence of a putative PBM at the extreme C-terminus (Fig. 1B). The Eukaryotic Linear Motif (ELM) resource (25) predicted that TF residues 71-76, with the sequence HGGSTV, constitute a class I PBM. This was confirmed by PDZPepInt, a PDZ domain identification software (26, 27), which further predicted that residues 71-75 (HGGST) can constitute a potential binding motif for the host protein Scribble. Mammalian Scribble, a cell polarity component with four PDZ domains, is targeted by multiple viral proteins through PDZ-PBM interactions (24). Particularly, domains 1 and 4 of Scribble have previously been reported to be high-affinity binders (28). We attempted to check the association of CHIKV TF with the PDZ binding domains of Scribble using *in silico* methods. A predicted structure of full-length TF (from the I-TASSER web server) was docked against PDZ domains 1, 2, 3, and 4 of Scribble (designated ScPDZ1, ScPDZ2, ScPDZ3, and ScPDZ4) using the ClusPro web server. The best-scoring models were selected for further analysis, and the binding interface was analyzed using Ligplot (Fig. 1C). MD simulations of TF in conjunction with ScPDZ1, ScPDZ2, ScPDZ3, and ScPDZ4 were subsequently carried out, with a PDZ domain from the mammalian High-temperature requirement A1 enzyme (HtrA1) (29) as a control. An RMSF plot (Fig. 1D) and RMSD (fig. S1) analysis of the simulation trajectory showed that the C-terminal tail of TF was stabilized by interaction with the Scribble PDZ domains. Local fluctuations in the TF C-terminus were significantly reduced upon interaction with PDZ domains 1 and 4 of Scribble, while the HtrA1 PDZ domain had minimal effect of TF stability (Table 1 and table S1). Specifically, TF-HtPDZ interaction was observed to be ∼3.8 times weaker than that with ScPDZ4 and ∼5.8 times weaker as compared to TF-ScPDZ1. The binding energies in all cases were primarily mediated by electrostatic and van der Waals interactions (Fig. 1C and table S1). The association of TF with ScPDZ2 and ScPDZ3 was also analyzed and the binding energies were found to be at least 2-fold lower than that of TF-ScPDZ1 and 1.4 times lower than that of TF-ScPDZ4 (Fig. S2 A, B and table S1,). Thus, *in silico* analysis indicated that CHIKV TF preferentially binds to the PDZ domains 1 and 4 of Scribble, as compared to PDZ domains from other mammalian proteins.

**Table 1.** Binding interface of docked complexes.

| System | CHIKV-TF<br>(Hydrogen bonds) | CHIKV-TF<br>(Hydrophobic interactions) |
| --- | --- | --- |
| TF-ScPDZ1 | Asp68, Glu70, Thr75 | Trp11, Trp19, Leu23, Ala27, His71, Gly72, Gly73, Ser74 |
| TF-ScPDZ4 | Glu70, His71, Gly72, Thr75 | Tyr3, Leu10, Trp11, Ala27, Ile30, Val31, Asn34, Leu38 |
| TF-HtPDZ | Gln15, Asn66, Glu70, His71, Gly73 | Leu10, Trp11, Leu20, Ser67, Asp68, Pro69, Gly72 |

### Overexpressed CHIKV TF co-localizes with Scribble

In order to investigate interactions between TF and Scribble biochemically, a plasmid containing the cDNA of CHIKV TF, with an N-terminal Enhanced Green Fluorescent Protein (EGFP) tag (EGFP-TF), was transfected in HEK293T cells. To check for the specificity of the putative interaction, the cDNA of CHIKV 6K, which shares the N-terminal forty-nine residues with TF, was similarly fused with EGFP to form EGFP-6K and transfected. The ribosomal slippage site was mutated in the 6K construct to prevent the initiation of TF expression from this plasmid (table S3). Twenty-four hours post-transfection, cells were fixed and immuno-stained with a Scribble-specific antibody to establish the intracellular localization of Scribble in the presence of CHIKV TF or 6K (Fig. 2A). In mock-transfected cells, Scribble was found to localize primarily to the cellular periphery near the plasma membrane. Similarly, Scribble localized to the plasma membrane in cells transfected with an empty vector (pEGFPC1), and no co-localization with EGFP was noted. However, in cells transfected with EGFP-TF, the localization of Scribble was drastically altered. In these cells, EGFP-TF and Scribble were found to co-localize in the form of distinct puncta in the cytoplasm (Fig. 2A), which was validated by line intensity plot analysis. No such co-localization was noted upon transfection of cells with EGFP-6K (Fig. 2A and fig. S 3). The line intensity profile graph displays the fluorescence signal intensity along the y-axis and the position or distance along the x-axis for two fluorochromes. The position of the intensity peaks indicated spatial overlap of TF and scribble proteins (Fig. 2A and fig. S3), which was further validated by Mander’s coefficient analysis (fig. S4). We hypothesized that the prominent punctation and mis-localization of Scribble is probably mediated by the unique C-terminal region of TF, since the N-terminal regions of 6K and TF are identical (Fig. 1A). Given the conclusions of the *in-silico* studies, we further hypothesized that the association between Scribble and TF is likely mediated by the PDZ domain(s) and PDZ-binding motifs of the components.

**Fig. 2.**
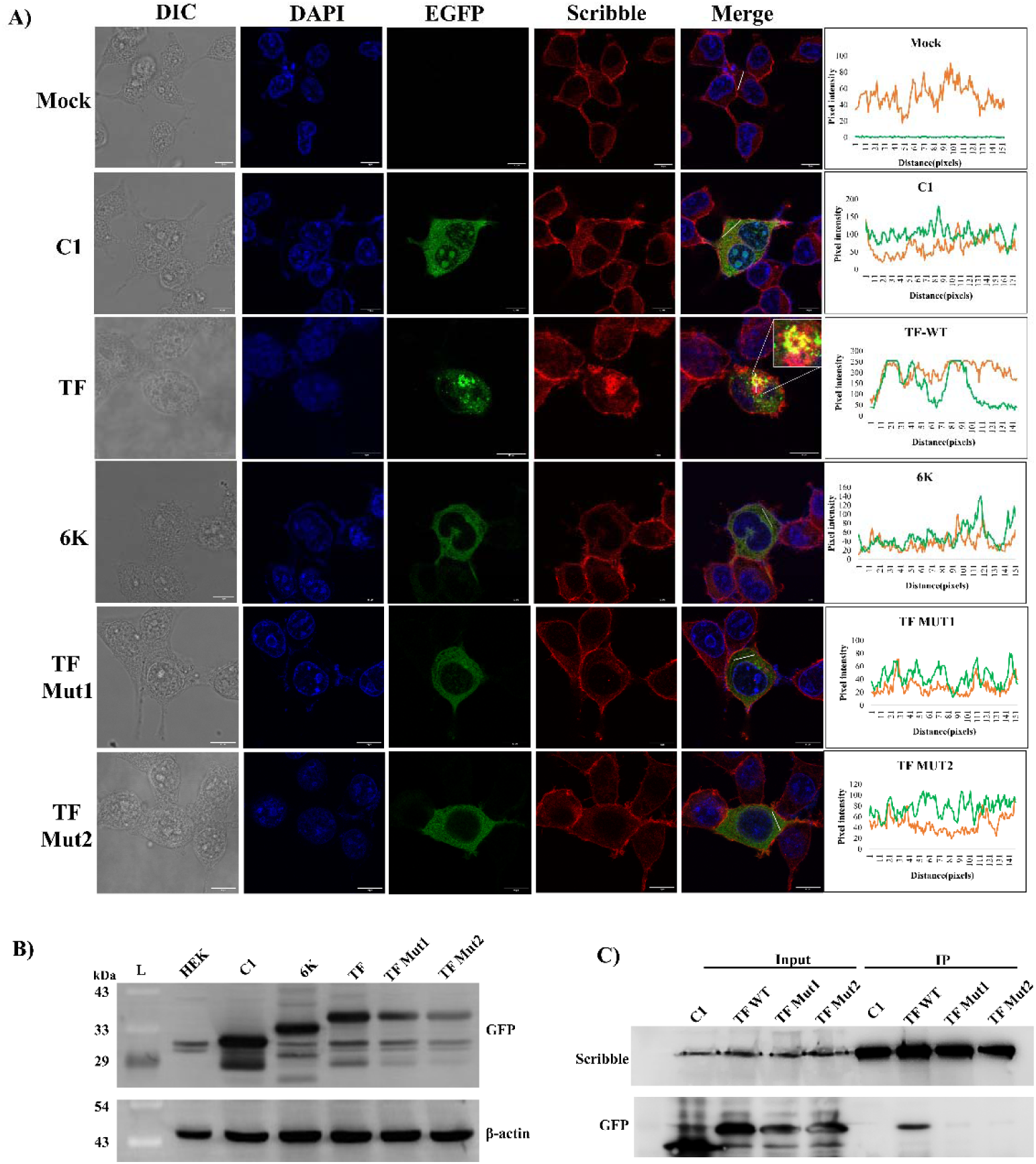
TF and Scribble association in transfected cells through confocal microscopy. HEK293T cells were transfected with pEGFPC1 vector or pEGFPC1 plasmids carrying TF or 6K. Twenty-four hours post-transfection, cells were fixed, and immunostaining was performed for Scribble (red). Inset shows zoomed images of an EGFP-TF expressing cell (A). Expression levels of TF and TF mutants in HEK293T cells. Twenty-four hours post-transfection, cells were lysed, and 6K and TF expression were analyzed using western blot with anti-GFP antibody. L: protein ladder, C1: pEGFPC1 empty vector, TF Mut1: EGFP-TF-HGAATV, TF Mut2: EGFP-TF-AAAATV (B). Immunoprecipitation (IP) of Scribble using Scribble specific antibody (i) and co-IP of TF-Scribble in transfected cells (C1, TF-WT, TF-Mut1 and TF-Mut2) (C). Fluorescent intensity profiles of the lined areas are displayed alongside the merged image in all cases. Scale - 10μm

### The interaction between TF and Scribble is mediated by a PDZ-PBM interaction

In order to investigate the association between TF and Scribble, the putative PBM in CHIKV TF (71-HGGSTV-76) was mutated to contain either two or four alanine substitutions (HGAATV/AAAATV). The constructs were designated TF Mut1 and TF Mut 2, respectively. The corresponding plasmids with the mutated EGFP-TF sequences were individually transfected in HEK293T cells, and the effect of these mutants on Scribble localization was analyzed using confocal microscopy.

We did not observe puncta formation by Scribble in cells transfected with the mutated EGFP-TF constructs (Fig. 2A). The fluorescence associated with TF Mut1 and TF Mut2 had a strikingly different distribution compared to wildtype TF. No co-localization was detected between Scribble and PBM mutated TF constructs, either visually or upon line intensity profile analysis (Fig. 2A and fig. S3). To confirm that the TF PBM mutants were being expressed in cells, the expression level of wildtype and mutated EGFP-TF constructs were investigated via Western blots (Fig. 2B). A slight reduction in the protein expression level of EGFP-TF with the PBM mutated to HGAATV (TF Mut1) compared to wildtype was observed, while the EGFP-TF construct with a quadruple alanine mutation (TF Mut2) displayed further reduced expression. However, the molecular weight difference between EGFP only, EGFP-6K and the EGFP-TF constructs clearly indicated the expression of the correct proteins in transfected cells (Fig. 2B). Further, alterations in the cellular expression patterns of the mutated constructs and the lack of co-localization with Scribble strongly suggested that the putative PDZ-binding motif HGGSTV is critical for the association between TF and Scribble (Fig. 2A).

To probe this interaction further, we immunoprecipitated Scribble with a rabbit polyclonal antibody from HEK293T cells transfected with EGFP only, and EGFP tagged TF, TFMut1 and TFMut2. The immunoprecipitated fractions were probed with an anti-Scribble antibody and an anti-GFP antibody to identify TF and TF mutants (Fig. 2C). We found that while TF could be immunoprecipitated with an anti-Scribble antibody; TFMut1 or TFMut2 could not be pulled down along with Scribble. This strongly indicates that the association between Scribble and TF is mediated via the PDZ domains of the former and the PBM of the latter, since mutating the PBM abrogated the association.

To establish the differences in Scribble binding by wildtype vs PBM mutated versions of TF, LC-MS/MS analysis of trypsin-digested fragments from the immunoprecipitated fractions was carried out. While wildtype EGFP tagged TF had a substantial coverage in the fractions immunoprecipitated by an anti-Scribble antibody, TF Mut1 and TF Mut2 had significantly reduced coverage and fewer peptides, less than half of that observed for the wildtype TF (Table 2), suggesting considerably lower association of the mutated versions of TF with Scribble. Taken together, co-localization studies, immunoprecipitation, and mass spectrometry clearly establish that the *in vivo* association between Scribble and CHIKV TF is driven by the interaction between the PDZ domain(s) of the former and the PBM at the unique C-terminus of TF.

**Table 2.** Mass spectrometric analysis of immunoprecipitate using anti-Scribble Ab.

| Sample | Accession | Description | Coverage [%] | # Peptides | # PSMs |
| --- | --- | --- | --- | --- | --- |
| **C1** | Q14160 | Protein scribble homolog<br>OS=Homo sapiens | 58 | 88 | 771 |
|  | GFP | GFP | - | - | - |
| **TF** | Q14160 | Protein scribble homolog<br>OS=Homo sapiens | 58 | 87 | 683 |
|  | TF WT | TF WT | 54 | 14 | 83 |
| **TF-MUT1** | Q14160 | Protein scribble homolog<br>OS=Homo sapiens | 58 | 94 | 721 |
|  | TF MUT1 | TF MUT1 | 28 | 7 | 38 |
| **TF-MUT2** | Q14160 | Protein scribble homolog<br>OS=Homo sapiens | 57 | 83 | 538 |
|  | TF MUT2 | TF MUT2 | 21 | 6 | 41 |

### TF associates with Scribble during CHIKV infection

In order to determine if the association of TF with Scribble has any functional relevance, the interaction was investigated during CHIKV infection of HEK293T cells. An antibody was raised against the unique region of CHIKV TF, by utilizing the C-terminal 27 amino acids coupled with Keyhole Limpet Hemocyanin (KLH) as an antigen. The antibody, generated in rabbit, was tested for its ability to recognize TF in western blots as well as in immunofluorescence assays (fig. S5). The association of the antibody with TF was found to be specific as it was able to recognize bacterially produced and purified TF (conjugated with the Maltose Binding Protein, MBP), as well as EGFP-TF from a HEK293T cell lysate, but was unable to bind to bacterially produced MBP-6K or overexpressed EGFP-6K (fig. S5).

The antibody against TF was utilized to visualize the localization and association of TF during CHIKV infection. Confocal microscopy studies showed substantial production of TF during CHIKV infection of HEK293T cells and TF co-localization with Scribble in punctate forms, as indicated by the line intensity profile analysis and Mander’s coefficient analysis (Fig. 3A and fig. S6).

**Fig. 3.**
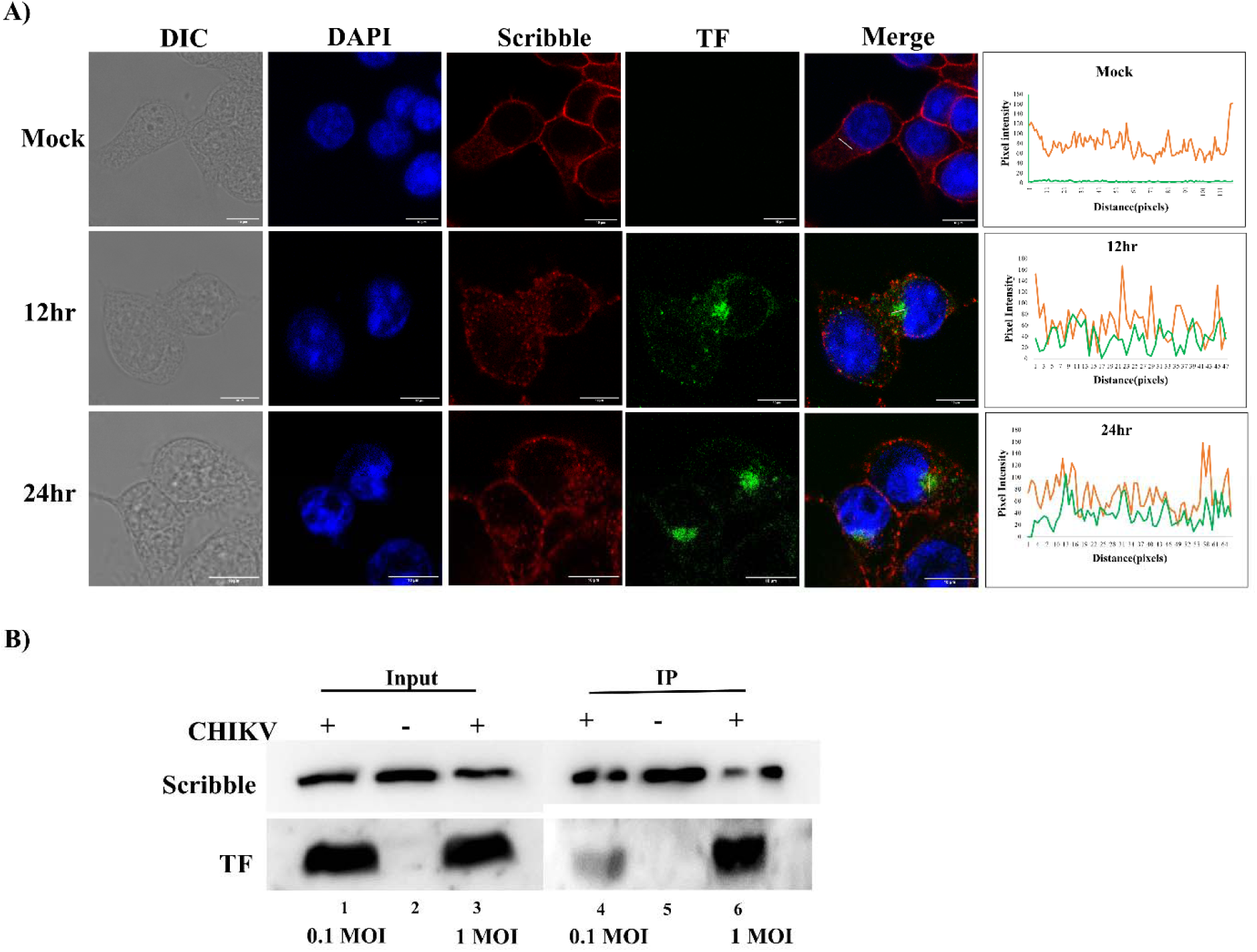
TF and Scribble association in CHIKV infected cells. HEK293T cells were infected with CHIKV at 0.1 MOI. Co-localization of TF-Scribble during infection was analyzed using TF and Scribble specific antibodies at the indicated time points (A), Immunoprecipitation (IP) of Scribble using Scribble specific antibody and co-IP of TF-Scribble in infected cells (B). Fluorescent intensity profiles of the lined areas are displayed alongside the merged image in all cases. Scale - 10μm.

The association between Scribble and TF during CHIKV infections was further established by immunoprecipitation of Scribble using a polyclonal anti-Scribble antibody from CHIKV-infected HEK293T cell lysates. Western blotting of the co-IP fractions with the TF unique-region specific antibody clearly indicated the presence of TF (Fig. 3B), thus establishing the association of TF with Scribble during CHIKV infection.

### CHIKV infection results in degradation of Scribble

Multiple viruses have been shown to target Scribble for degradation in order to manipulate host immune responses, cell junction integrity, and apoptosis pathways (24). In order to analyze the effect of CHIKV infection on the level of Scribble, the levels of Scribble protein were detected by western blotting in HEK293T cells 24 h and 48 h post infection (Fig. 4A). A significant decrease in Scribble protein levels was observed following CHIKV infection in a time-dependent manner. At 24 hours post-infection, Scribble levels were reduced to nearly 50% compared to uninfected cells. The protein levels reduced to greater than 95% at 48 h post infection (Fig. 4A), while those in uninfected cells remained at the initial level. Thus, CHIKV infection appeared to cause a drastic reduction in the levels of human Scribble protein in cell culture.

**Fig. 4.**
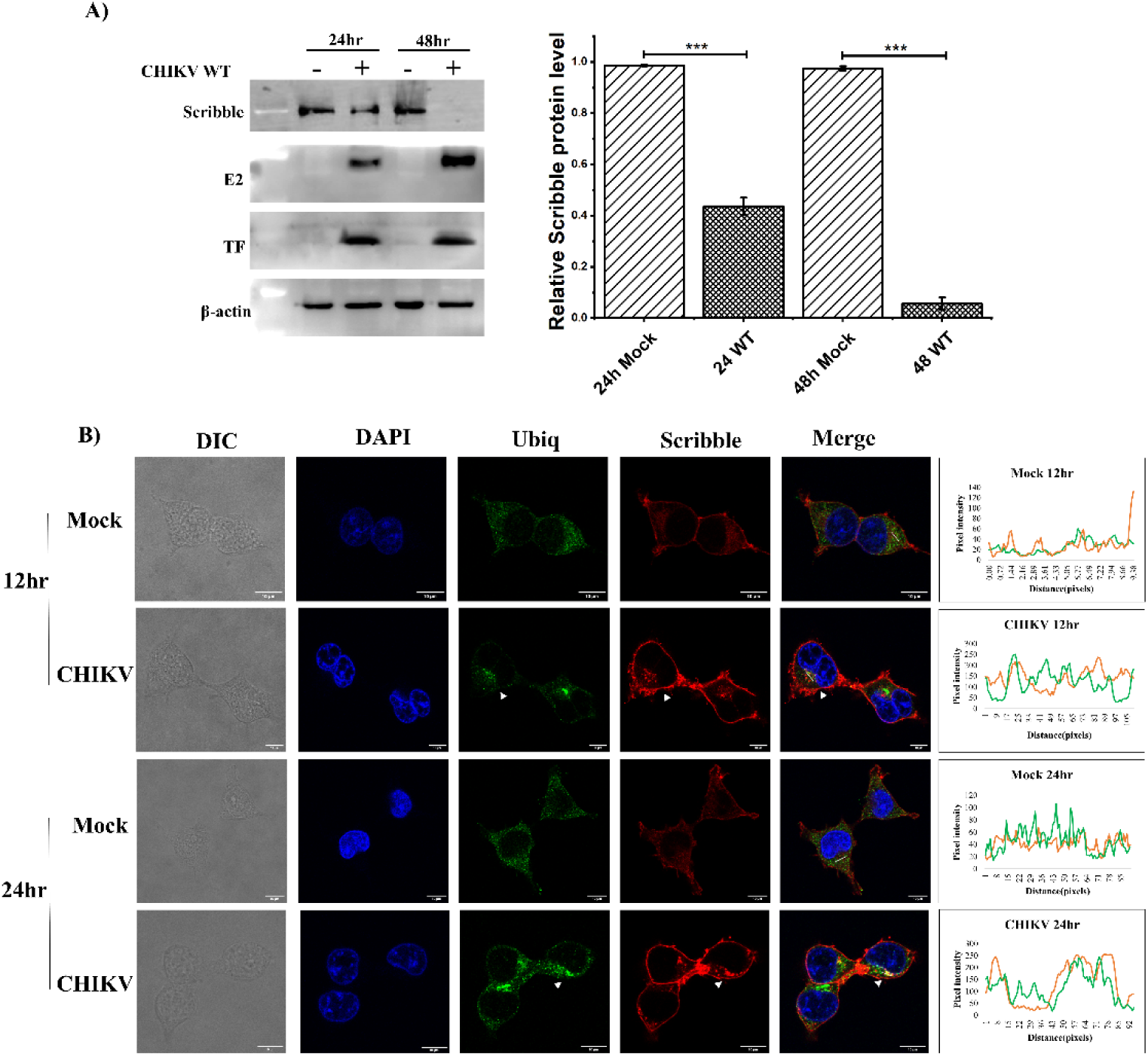
Effect of CHIKV infection on Scribble localization and expression. HEK293T cells were infected with CHIKV and Scribble levels were analyzed at 24 and 48 hpi using western blot and quantitated using densitometric analysis (right panel) (A), Immunostaining for Ubiquitin (Ubiq) and Scribble in CHIKV infected HEK293T cells at the indicated time point post-infection. Arrowheads (white) indicate Scribble puncta and co-localization with ubiquitin (B). Data represent mean ± SEM. **, *p* ≤ 0.01; \*\**\**, *p* ≤ 0.001, ns: non-significant (Student’s *t-*test). Fluorescent intensity profiles of the lined areas are displayed alongside the merged image in all cases. Scale - 10μm.

### Scribble degradation during CHIKV infection is through the Ubiquitin-Proteasome System

The degradation pathway of Scribble was explored by co-staining CHIKV-infected cells with anti-Scribble and anti-Ubiquitin primary antibodies, and appropriate secondary antibodies (Fig. 4B). We found co-localization of the Scribble puncta with ubiquitin-specific antibodies in CHIKV-infected HEK293T cells, which suggests that CHIKV-induced degradation of Scribble could be mediated by the Ubiquitin-Proteasome System (UPS).

Similarly, in transfected cells overexpressing TF, co-localization of Scribble, Ubiquitin, and TF was detected in cytoplasmic puncta (Fig. 5A). While some Scribble punctation, albeit at a lower level, was noted during overexpression of the double alanine PBM mutant (EGFP-TF-HGAATV), there was no co-localization of ubiquitin to these puncta (Fig. 5A and fig. S7).

**Fig. 5.**
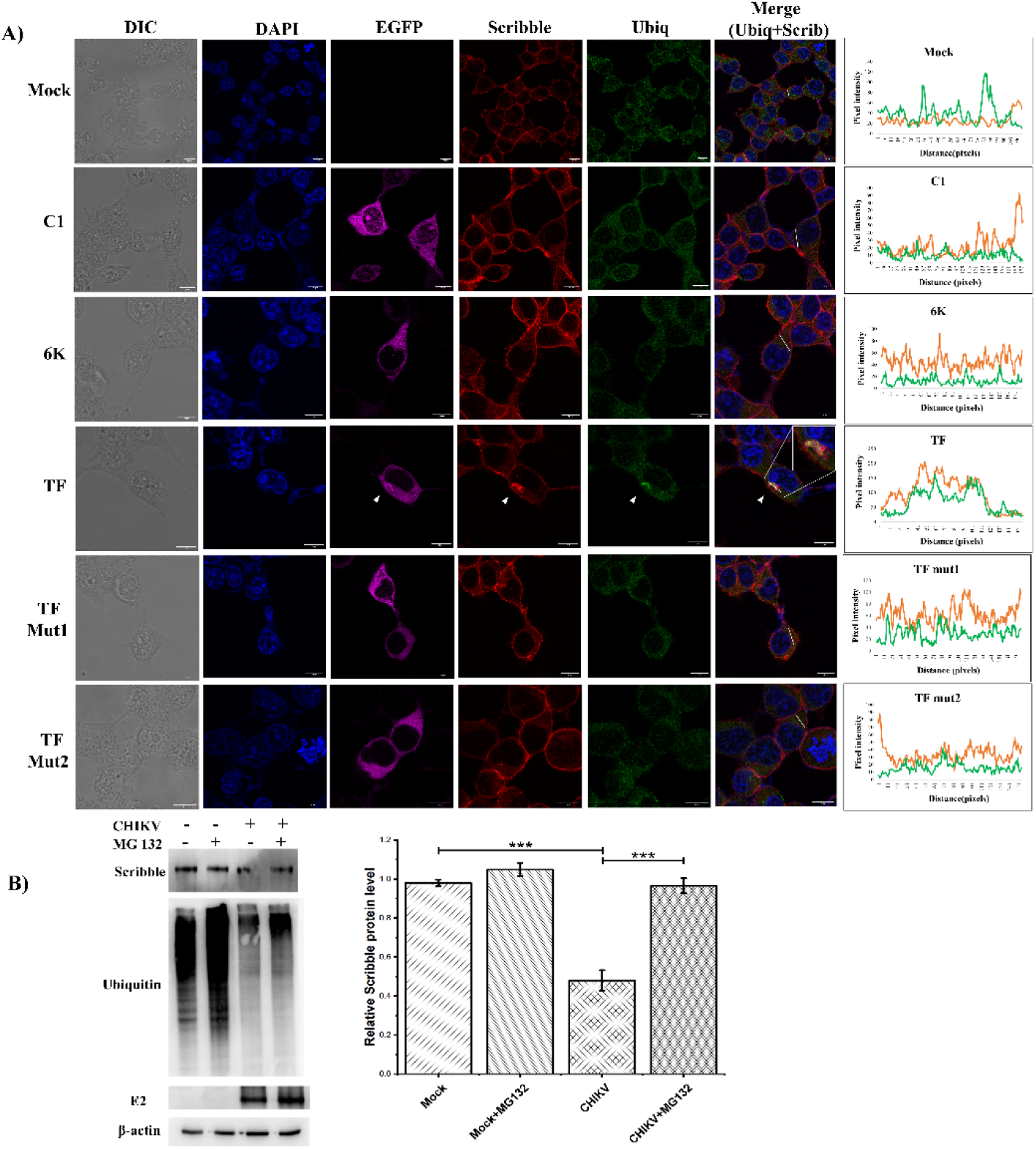
Effect of TF on Scribble ubiquitination. HEK293T cells were transfected with pEGFPC1 vector or pEGFPC1 plasmids carrying TF or 6K. Twenty-four hours post-transfection, cells were fixed, and immunostaining was performed for Scribble (red) and Ubiquitin (green). Inset shows a zoomed image of an EGFP-TF expressing cell. Arrowhead (white) indicates EGFP-TF, Scribble, and ubiquitin puncta and co-localization in the merged image (A). HEK293T cells were infected with CHIKV, treated with MG132, and Scribble levels were analyzed at 24 hpi using western blot (B). Data represents mean ± SEM. **, *p* ≤ 0.01; \*\**\**, *p* ≤ 0.001, ns: non-significant (Student’s *t-*test) (C). Fluorescent intensity profiles of the lined areas are displayed alongside the merged image in all cases. Scale - 10μm.

To further investigate the involvement of the UPS in Scribble degradation, ubiquitinated proteins in CHIKV infected or uninfected HEK293T cells were analyzed in the presence of the proteasome inhibitor MG-132 (Fig. 5B). In uninfected cells, the addition of MG-132 had no effect on Scribble quantities. However, the addition of MG-132 restored the reduction in Scribble triggered by CHIKV infection.

Thus, taken together, our results suggest that during CHIKV infection, the TF component associates with Scribble through its C-terminal PBM (HGGSTV), resulting in the punctation and degradation of Scribble. The alterations in Scribble quantities upon the addition of MG132 suggest that the degradation of Scribble probably occurs through the UPS pathway.

### Scribble knockdown enhances CHIKV infection

In order to investigate the significance of Scribble downregulation during CHIKV infection, the level of CHIKV propagation in Scribble knockdown cells was compared to that in untreated cells. Two short hairpin RNAs (shRNAs) were used for the knockdown of Scribble protein expression in HEK293T cells according to standard protocols. The effect of the shRNAs was analyzed 36 hours post-transfection using western blot analysis. Densitometric analysis, to quantitate the relative percentage of gene knockdown for each sample, indicated a relatively efficient Scribble protein knockdown of approximately 50% with both shRNA conditions (fig. S8 A). The degree of knockdown at the RNA level was also confirmed using quantitative real-time polymerase chain reaction (qRT-PCR) (fig. S8 B). To ensure that any alteration observed in CHIKV infectivity upon Scribble knockdown was indeed because of a reduction in the quantity of Scribble, and not due to a difference in the number of viable cells as a result of knockdown, cell viability was assessed using an MTT assay (fig. S8 C). No significant change in cell viability was observed with Scribble knockdown.

The HEK293T cells with reduced Scribble expression were then infected with CHIKV (strain 119067) at an MOI of 0.1. The culture supernatants containing the new virus were harvested 24 h post infection and a plaque assay was performed using standard methodologies in Vero cells. CHIKV infectivity was calculated by counting the number of plaques formed 48 h post-virus addition. We found that the virus titre, represented as the plaque-forming unit per ml (pfu/ml), was significantly increased upon Scribble knockdown (Fig. 6A). We also tested the levels of CHIKV RNA in infected HEK293T cells with Scribble knockdown by RT-PCR and found that the levels of CHIKV RNA were also markedly upregulated in these cells, compared to regular infected cells (Fig. 6B). The increase in infectious CHIKV virions and genomic RNA was approximately 2-3 times higher upon Scribble knockdown. Taken together, our data strongly suggest that punctation and degradation of Scribble is a viral countermeasure to facilitate CHIKV propagation; and that this activity is mediated by TF.

**Fig. 6.**
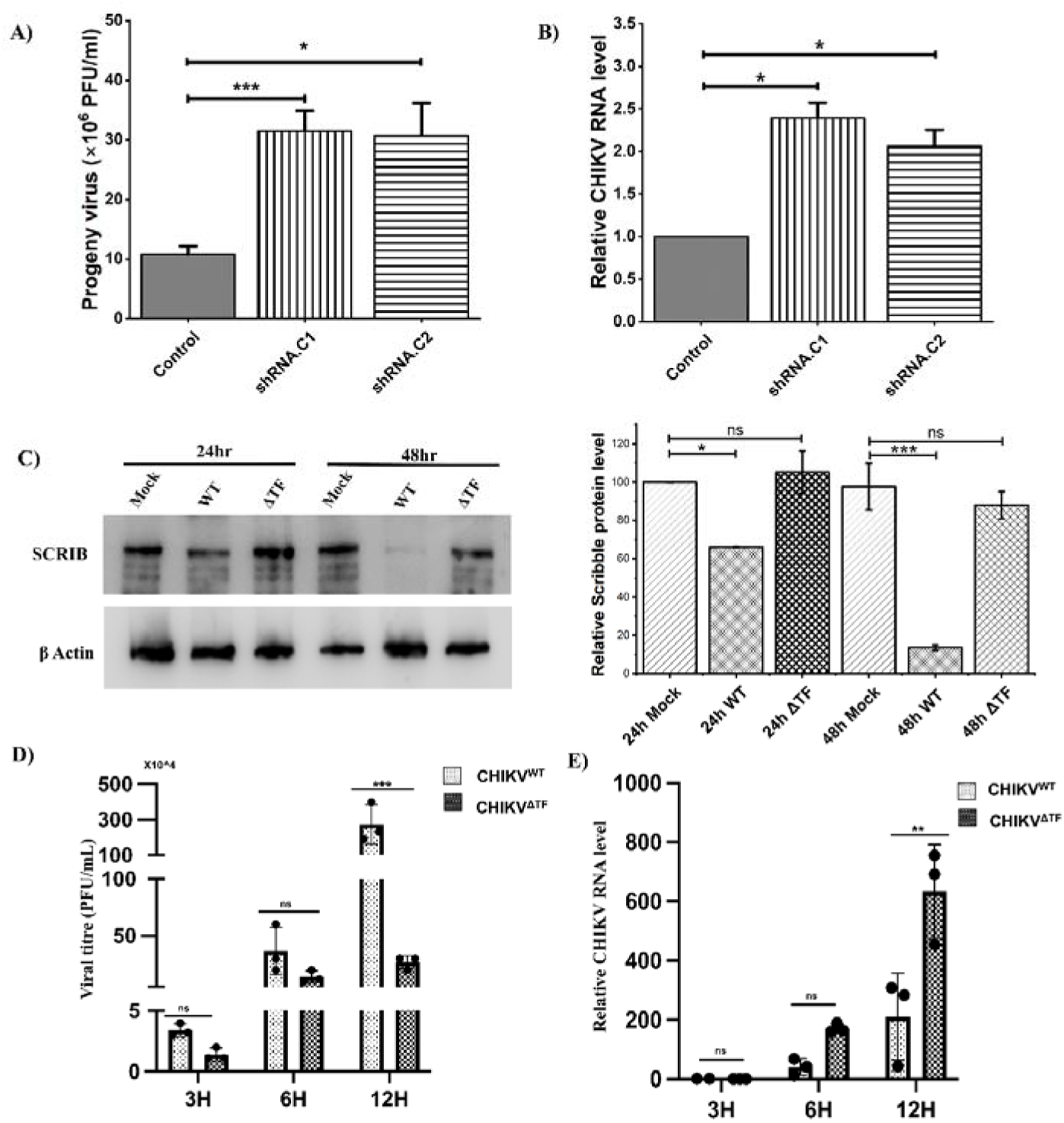
TF-deficient CHIKV exhibits reduced cellular propagation, and Scribble knockdown modulates CHIKV infection. Scribble shRNAs were transfected into HEK293T cells, thirty-six hours post-transfection, cells were infected with CHIKV at MOI of 0.1 for 24 hours. The effect of Scribble knockdown on CHIKV progeny virus was determined using plaque assay (A) and CHIKV RNA levels using real-time PCR (B). Western blot depicting scribble expression in HEK 293T cells infected with 0.1 MOI WT mcherry CHIKV and ΔTF mcherry CHIKV at 24 hours post-infection and 48 hours post-infection and densitometric analysis (right panel) (C). The infectivity of WT and ΔTF CHIKV compared using plaque assay (D), and by measuring respective RNA levels using real-time PCR (E). Data are presented as mean ± SEM (*n* = 3). Statistical analysis was performed using an unpaired two-tailed Student’s *t*-test for two-group comparisons and one-way ANOVA with Tukey’s post-hoc test for multiple-group comparisons. *P* < 0.05 was considered statistically significant.

### CHIKV lacking TF is compromised in cellular propagation

To further investigate the role of TF during CHIKV infection, a mutated version of CHIKV was generated to produce all structural proteins including 6K, but not TF. An infectious clone of the CHIKV OPY-1 strain with an mCherry cassette for detection (mCherry-LR2006-OPY1) (30) was utilized as a template to generate a silent mutation in the ribosomal slippage site, to abrogate the production of TF (fig. S9). Both infectious clones WT CHIKV and ΔTF CHIKV, were transfected in HEK293T cells using the standard protocol for the formation of virus particles. The transfection supernatants were subjected to plaque assays and the viral plaques in each case were isolated and subjected to serial infections to increase the titre. We confirmed, using specific antibodies, that while infection with WT CHIKV results in the production of TF as well as other structural proteins; infection with ΔTF CHIKV produces 6K, but not TF (fig. S10).

Cellular lysates from WT and ΔTF CHIKV OPY-1 infected cells were subjected to western blotting with anti-Scribble antibody at 24 hours p.i. and 48 hours p.i. A significant, time-dependent decrease in Scribble protein levels was observed in cells infected with the WT CHIKV OPY-1 strain. In contrast, cells infected with ΔTF CHIKV OPY-1 showed Scribble levels slightly above normal (Fig. 6C). Together, these results demonstrate that the TF component of CHIKV is crucial for effective downregulation of Scribble, a process that cannot be executed by ΔTF CHIKV.

Quantification of infectious virus particles from the media was carried out for both WT and ΔTF CHIKV. The plaque assay consistently suggested lower infectious particle numbers for ΔTF CHIKV compared to WT virus (Fig. 6D). However, qRT-PCR based quantification of viral RNA for WT and ΔTF CHIKV showed higher viral RNA load in cells infected with ΔTF CHIKV 12 hpi (Fig. 6E). This indicates production of significantly higher numbers of progeny virions that are defective in release or cellular propagation when TF is absent.

To explain this observation on lack of efficient propagation of ΔTF CHIKV, the cellular boundaries in the case of WT and ΔTF CHIKV infected cells were observed by antibody-mediated staining of β-catenin, a key adhesion protein that associates with E-cadherin (31). A drastic difference in β-catenin staining, indicating major alterations in cellular boundaries, wa observed in WT CHIKV infected cells, while the morphology of cell-cell association regions in ΔTF CHIKV infected cells was similar to that of uninfected cells (Fig. 7). This drastic alteration in the morphology of cellular boundaries in WT CHIKV suggests easier routes for escape and propagation for infectious WT particles, which cannot be effectively utilized by ΔTF CHIKV. Taken together, these observations emphasize the requirement for cellular polarity/adhesion disruption mediated by the mislocalization and degradation of Scribble by TF, for optimal CHIKV transmission *in vitro*.

**Fig. 7.**
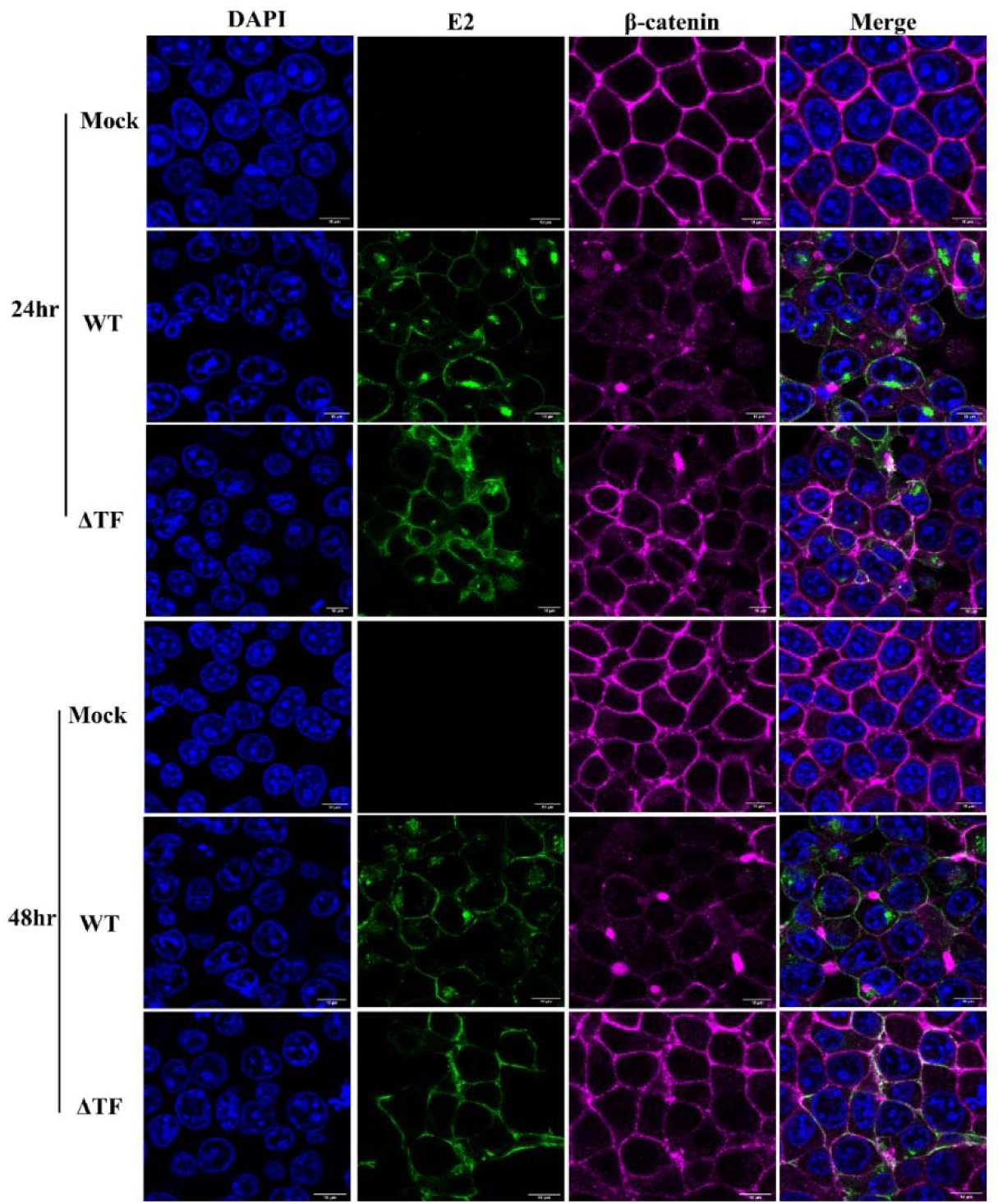
TF-deficient CHIKV alters cell-cell boundaries. HEK293T cells were infected with WT CHIKV and ΔTF CHIKV at 0.1 MOI. Cells were then stained for β-catenin and E2 at 24 hpi and 48 hpi. Scale - 10μm.

### A cryoEM reconstruction of **Δ**TF CHIKV indicates **Δ**TF CHIKV is only impeded in terms of cellular egress, and not in entry or assembly

In order to detect any structural alterations in the ΔTF CHIKV, the particles were purified, cryofrozen and single particle reconstruction was carried out from cryoEM data (Fig. 8). 2D and 3D classification indicated the presence of one predominant species (fig. S12). The final reconstruction was carried out with 84,104 particles with icosahedral symmetry imposed and a map at a global resolution of 3.6 Å (EMD-69164) was generated. A reconstruction of wild-type CHIKV particles treated similarly yielded a map with a global resolution of 4.31 Å (EMD-69185) (Fig. 8A-F).

**Fig 8.**
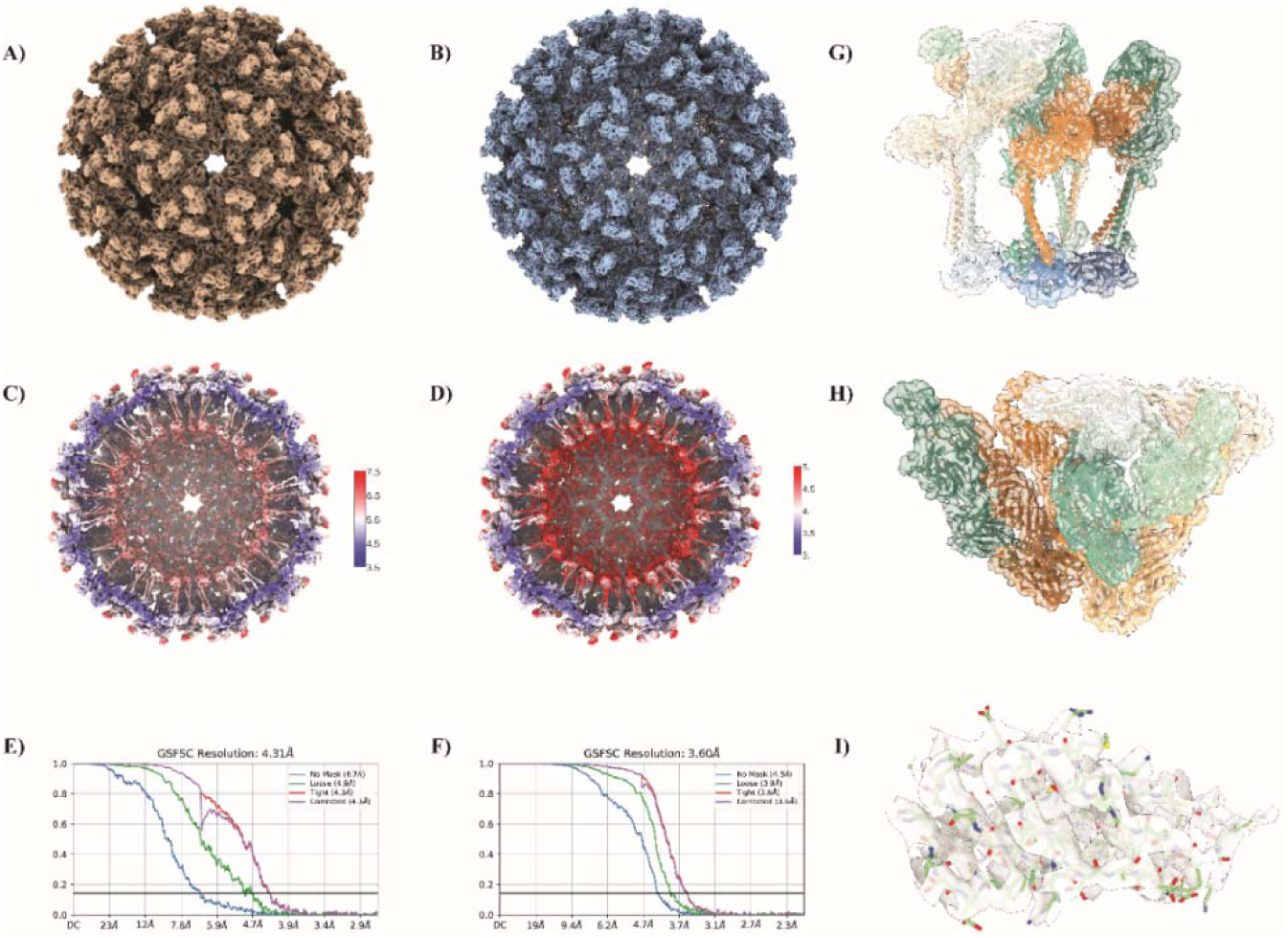
Cryo-EM maps and validation of the CHIKV structures. Density map for the wild-type (wt) CHIKV(A). Density map for ΔTF CHIKV (B) . Corresponding local resolution section maps for the structures in (A) and (B), respectively (C) and (D). Gold-standard Fourier Shell Correlation (FSC) curves used for resolution estimation of the CHIKV WT and ΔTF maps respectively (E) and (F). PDB model of the CHIKV asymmetric unit (8FCG) fitted into the ΔTF CHIKV asymmetric unit (ASU) density, displayed as side and top views carved at 5 Å. Capsid monomers are shown in shades of blue (pale sky blue to deep navy), E1 monomers in shades of orange (pale apricot to deep ochre), and E2 monomers in shades of green (pale mint to deep pine)(G) and (H). A focused view of the E1 protein domain II - β barrel region, residues 308–370, carved at 2 Å using the sharpened map(I).

An initial analysis of the map revealed the traditional double layered particle structure of alphaviruses, with the outer glycoprotein layer consisting of E1-E2 heterodimers and the inner layer of nucleocapsid proteins, arrayed in accordance with T=4 icosahedral symmetry. The map for ΔTF CHIKV is at a global resolution of 3.6 Å, which is among the highest resolution alphavirus cryoEM structures reported to date (32). The model for one icosahedral asymmetric unit fitted well in the segmented density for one iASU of ΔTF CHIKV (Fig. 8G-I), indicating that particle assembly is accurate, and all other structural components are correctly placed in the TF deficient version of CHIKV. As there are no alterations in the positioning of the E1-E2 glycoproteins, it can be conjectured that ΔTF CHIKV is capable of receptor binding and entry through the usual route (33). Therefore, the defect in the context of ΔTF CHIKV transmission appears to be entirely associated with the lack of TF, and its interaction with Scribble for modification of cellular boundaries.

## Discussion

Host-virus interactions play a crucial role in defining the trajectory of pathogenesis (34). Key host molecules and pathways are consistently hijacked by viruses to ensure the propagation of infection. Some widely targeted host cellular co-factors include nuclear pore/envelope complexes, COPI vesicles, ribosomes, and the translational pathway and signalling modulators (35). Increasing evidence suggests that the Scribble cell polarity complex is a central host factor targeted by multiple virus families (24). Human Scribble is part of a large module of adapters, including several Dlg and Lgl proteins, which has a central role in regulating cell migration, adhesion, polarity, signalling, apoptosis, morphogenesis, and development (36–38). Members of this complex associate with a wide range of cellular factors to modulate cellular activities. Scribble is a member of the LAP (leucine-rich repeats and PDZ domain) family and contains numerous protein-protein interaction domains, including 16 Leucine-Rich Repeat domains (LRRs), 2 LAP-specific domains, and 4 PDZ domains (PDZ1-4). Loss of Scribble disrupts cellular adhesion and cell polarity, leading to hyperproliferation (36). As a determinant of cellular growth, apoptosis and proliferation, Scribble has been a target of tumorigenic viruses such as Human T-lymphotropic virus 1 (HTLV-1) and Human Papillomavirus (HPV) (39,40). Recently, effectors from several other viruses, such as the NS1 protein from Influenza A Virus (IAV), NS5 from Tick-borne encephalitis virus (TBEV), and the core protein from Hepatitis C

Virus (HCV), have been shown to associate with the PDZ domains of Scribble through PBMs, resulting in mistargeting of Scribble and an increase in viral proliferation (24). However, there have been no reports of Scribble modulation by alphaviruses so far. In this manuscript, we establish Scribble as a host target of CHIKV and identify the viral factor that induces Scribble mislocalization and degradation, leading to major structural alterations at cellular boundaries to enhance virus release and proliferation.

We found that CHIKV infection downregulates Scribble expression by as much as 95% in cultured cells in a time-dependent manner. Immunofluorescence analysis of CHIKV-infected cells showed clustering of Scribble into cytoplasmic puncta and ubiquitin-mediated degradation. Degradation of Scribble via the Ubiquitin-Proteasome pathway was suggested by experimental studies using a proteasome inhibitor, MG132. The cellular localization of Scribble is vital for its functioning (24, 22, 38), and mislocalization of Scribble and its sequestration in discrete puncta likely affect its regulatory role in cellular proliferation (39, 41). To confirm the role of Scribble in CHIKV infection, shRNA-based downregulation of Scribble expression was carried out. A ∼50% downregulation in Scribble levels resulted in 2-3 times increase in the virus titer and genomic RNA levels. Scribble is a cell polarity protein that is important for maintaining the integrity of cellular junctions, and disruption of cell junctions has been reported to promote virus egress and dissemination (23,24,36,42). While HEK293T cells do not contain conventional tight junctions, they do express a number of cell adhesion molecules, including E-cadherin, occludin, and ZO-1, in addition to Scribble (43). Scribble is known to associate directly with E-cadherin and β-catenin to maintain the integrity of cellular junctions (44–46). Thus, the beneficial effect of Scribble knockdown on CHIKV replication is likely due to altered viral release.

6K and its transframe variant, TF, play a significant role in regulating CHIKV morphology and virus-host interactions (47). The ion channel activity of CHIKV 6K has been established (10). Inhibition of 6K activity disrupts CHIKV assembly, leading to the formation of morphologically unusual particles with reduced infectivity. Recent work has shown that 6K co-localizes with alphavirus glycoproteins in cytopathic vesicles (CPV-II), which may support its role in promoting viral assembly (14). TF, in contrast, localizes at the plasma membrane and is packaged in virions (15, 17). While TF can also form ion channels *in vitro*, the predicted structure of a multimeric TF channel is wider and less intricate, potentially offering reduced control over ion passage compared to 6K channels (19), suggesting that the primary function of TF lies elsewhere. Our work establishes a critical role for TF in modulating cellular boundaries and enhancing viral proliferation. We show conclusively that CHIKV TF is primarily responsible for downregulating Scribble, leading to alterations in cell-cell boundaries. Overexpression of TF in HEK293T cells leads to the sequestration of Scribble into distinct puncta, ubiquitination, and degradation, similar to that in CHIKV infected cells. Sequestration and ubiquitination of Scribble are not evident in cells transfected with CHIKV 6K, suggesting that TF, and not 6K, is the regulator of Scribble. TF can be immunoprecipitated with an anti-Scribble antibody from cells infected with WT CHIKV, indicating a physical association between the two components. Infection with ΔTF CHIKV resulted in substantially enhanced viral RNA load, coupled with reduced infectious particles in the supernatant, indicating that the lack of TF indeed prevents the release and propagation of CHIKV. β-catenin staining of cells infected with WT CHIKV showed clear disruption at the cellular boundaries. In contrast, ΔTF CHIKV-infected cells showed a similar pattern of β-catenin staining to uninfected cells, suggesting that the effective release of virus particles requires disruption of cellular boundaries and highlighting the critical role of TF in enhancing virus escape. Since TF and 6K share approximately two-thirds of their amino acid sequences, this functional difference is likely to originate in the unique C-terminal region of TF, which contains the PDZ domain-binding motif (PBM) as a prominent feature. The homologues of CHIKV TF from several other alphaviruses also have PBMs in their unique C-terminal tails that can potentially associate with host partners (fig. S11, table s2).

A high resolution cryoEM reconstruction of ΔTF CHIKV clearly indicated correct positioning and orientation of glycoproteins, suggesting that particle assembly is accurate and particles are not likely to be entry defective. The visual distinction between the boundaries of WT and ΔTF CHIKV infected cells suggest that the egress of ΔTF CHIKV may follow an atypical pathway compared to other alphaviruses. Further, while TF is thought to be a structural component of alphaviruses (18), no specific density for TF was noted in our WT CHIKV reconstruction. Both reconstructions were carried out with icosahedral symmetry imposed, which may result in averaging out of densities present in non-stoichiometric quantities. It is also possible that TF is primarily a non-structural component involved in host partner interaction, while a subpopulation may be associated with virus particles in non-stoichiometric fashion. Thus, asymmetric reconstructions at high resolution may be required to understand subtle, local differences in the morphologies of WT and ΔTF CHIKV, if any; and to identify the location of TF, if present, in WT alphavirus particles.

Other viral proteins, such as IAV NS1, TBEV NS5, HTLV tax, HCV NS4B, and HPV-16 E6 are known to associate with the PDZ domains of Scribble through their PBMs (24). *In silico* analysis of the unique C-terminus of TF indicated the presence of a Class I PBM which could interact with the PDZ domains of Scribble. Mutational analysis of the predicted PBM confirmed its involvement in regulating Scribble localization. The TF constructs with the PBM mutated were mislocalized, distributed throughout the cytoplasm, and unable to co-localize with or trigger the sequestration and ubiquitination of Scribble. Mutations in TF PBM also resulted in somewhat decreased protein levels, particularly in the quadruple alanine mutant, which could be due to the drastic nature of the mutation. However, the expressed protein showed an altered localization pattern distinct from that of the wild-type TF. Further, while GFP-TF with an intact PBM could be immunoprecipitated from transfected cells, TF constructs with mutated PBMs could not be pulled down with an anti-Scribble antibody. MS analysis confirmed the weakening of Scribble-TF association upon mutating the TF PBM, which suggests a central role for the PDZ-PBM interaction in this association. Interestingly, the association of Scribble with β-catenin, which is required for structural maintenance of cellular adhesion and boundaries, depends on the direct interaction between a PBM of β-catenin with the PDZ domains in Scribble (44–46). A competitive dissociation of β-catenin PBM binding by Scribble, in favour of TF PBM binding, may result in punctation and cytoplasmic mis-localization of Scribble, and consequent distortion of cellular boundaries as seen in cells infected with WT CHIKV. A loss in the β-catenin population in CHIKV infected cells was reported earlier and attributed to the non-structural nsp2 protein of CHIKV, however, this hypothesis was not tested in the context of a CHIKV mutant lacking nsp2 (48).

We note that while transfection vs viral infection-based production of TF had similar effects on Scribble; the reduction in Scribble protein quantity was marginal in the former case compared to the latter. This could be because of temporal alterations in TF expression in the transfection versus the infection model, or due to the differences in the quantities of TF production. Despite the high transfection efficiency of HEK293T cells, the expression efficiency of EGFP-TF or EGFP-6K by transfection was consistently low, which could be due to the membrane permeabilization capability of these components. Expression of 6K in bacterial cells impedes the growth curve significantly, as the membrane activity is detrimental to the cells (11,12). Thus, reduced levels of TF expression in transfected cells may be a possible reason for decreased levels of Scribble degradation, compared to those observed in CHIKV-infected cells. However, the decline in Scribble protein levels triggered by transfection of EGFP-TF was absent in the case of PBM-mutated TF constructs, which suggests the crucial role of TF PBM in PDZ-domain dependent association with Scribble.

A lingering question centres on how punctation of Scribble via TF may lead to its degradation. Other viral proteins, such as the Envelope protein (E) of betacoronaviruses and E6 of Papillomaviruses are known to associate with cellular junction proteins through PDZ-PBM binding and trigger their degradation, resulting in destabilization of the cellular boundaries. TF has the ability to form ion-channels in membranes, and therefore can be categorized as a viroporin (19), like the betacoronavirus E protein (49, 50). It will be interesting to determine whether, in either case, the viroporin activity is mechanistically related to the degradation of proteins associated with cell polarity or cellular junctions. Sequentially dissimilar viroporins encoded by diverse virus families may be performing analogous functions to ensure viral propagation across damaged cellular boundaries. Mechanistic analysis of this targeted junctional activity will require detailed future studies, and may necessitate an understanding of the cooperativity, if any, between the PBM and viroporin regions, and structure guided attenuation of viroporin function within TF.

In summary, we establish that the manipulation of host cell morphology by alphavirus TF is an adaptive strategy that increases the efficiency of pathogen propagation. The previously enigmatic TF protein is shown to play an important role in facilitating virus egress by targeting the cellular polarity complex. Our work suggests that alphavirus TF can be a druggable target for effectively reducing viral replication and generating pan-alphaviral inhibitors.

## Materials and methods

### Docking and Molecular Dynamics (MD) simulation

Since there is no experimental structure available for CHIKV TF, a structure predicted using the I-TASSER web server (51, 52) was utilized for docking. The model with the highest c-score was selected and further refined using the GalaxyRefine web server (53–55). The X-ray crystallographic structures of Scribble PDZ domains (1-4, referred to as ScPDZ1, ScPDZ2, ScPDZ3, and ScPDZ4) and mammalian HTRA1 PDZ (HtPDZ) domain were downloaded from the Protein Data Bank (PDB IDs: 5VWC, 1UJU, 7JO7, 4WYT and 2JOA respectively). All structures were energy minimized prior to docking and simulation studies.

Protein-protein docking between TF and the PDZ domains of Scribble and HTRA1 was carried out using the ClusPro web server (56–58). The binding interface and affinity of each docked complex were analyzed using LigPlot (59). All-atom MD simulation was carried out for each complex in triplicates as described. Briefly, GROMACS (60) was used to set up and execute the simulations. Each complex was solvated separately using the SPC water model (61) and neutralized using counter-ions. Periodic boundary condition was applied to each system. Energy minimization was executed using the steepest descent algorithm (62). Position-restrained NVT-NPT ensemble equilibrations were carried out for 500 ps and 1 ns respectively. Finally, the systems were subjected to unrestrained dynamics for 100 ns. Nose-Hoover (63) and Parrinello-Rehmann (64) were used for maintaining the temperature (300K) and pressure (1 bar) during simulations. Electrostatics was maintained using Reaction-Field (65) and LINCS (66) was used as the constraints algorithm. A time-step of 2 femtoseconds and the Verlet cutoff scheme was applied. All simulations were carried out using the Gromos 54a7 force field. Trajectory analysis was carried out using the built-in GROMACS tools. VMD (67) and UCSF Chimera (68) were used as the visualization tools.

### Molecular Mechanics Poisson-Boltzmann Surface Area (MM-PBSA) analysis

MM-PBSA was utilized to determine the Gibb’s free energy (GFE) of protein-protein interactions. The script g_mmpbsa (69) was used for evaluating the GFE of TF and PDZ domains from simulation trajectories. The following equations were used for the analysis

ΔG *bind* = ΔG *complex* – (ΔG *protein1* + ΔG *protein2*)

ΔGX = ΔEMM + ΔG *solv* - TΔSMM

ΔEMM = ΔE *bonded* + ΔE *nonbonded* = ΔE *bonded* + (ΔE *elec* + ΔE *vdw*)

ΔG *solv* = ΔG *polar* + ΔG *nonpolar*

> where Δ*G complex* = total free energy of protein-protein complex

> Δ*G protein1*, Δ*G protein2* = free energies of proteins in solvent

> Δ*GX* = free energy for each entity

> Δ*E MM* = vacuum potential energy (combined bonded and nonbonded interactions)

> Δ*G solv* = solvation free energy [polar (Δ*G polar*), nonpolar (Δ*G nonpolar*)]

> *T*Δ*S* = entropic contribution of free energy in vacuum. T is temperature and ΔS is entropy respectively.

TΔS MM was not included in this study as g_mmpbsa does not calculate entropy (S). Hence, ΔG *bind* was calculated as follows,

ΔG *bind* = (ΔE *elec* + ΔE *vdw*) + (ΔG *polar* + ΔG *SASA*)

where, ΔG *SASA* = ΔG *nonpolar* since nonpolar solvation energy was calculated based on the SASA model in this study.

### Cells and plasmids

Human embryonic kidney (HEK) 293T and Vero cells were grown in Dulbecco’s modified Eagle medium (DMEM) (Gibco) supplemented with 10 % fetal bovine serum (FBS) (Gibco) and 1 % antibiotic antimycotic solution (Himedia). The cells were maintained at 37 °C with 5 % CO_2_. The cDNA sequences of CHIKV 6K and TF (fig. S8) were cloned in the pEGFPC1 vector (Clontech) to generate fusion proteins tagged with the Enhanced Green Fluorescent Protein (EGFP) at the N-terminus. The ribosomal slippery site UUUUUUA in the 6K cDNA sequence was silently mutated to UUUCCTG to disrupt the consensus ribosomal frameshifting sequence and produce only 6K without affecting its primary structure (15). Alanine substitution mutations were carried out to disrupt the PDZ domain binding motif within amino acids 71-74 of TF. Two mutants – a) G73A S74A and b) H71A G72A G73A S74A, were generated using sequential site-directed mutagenesis (Life Technologies) and were confirmed by DNA sequencing (table S3). The ribosomal slippery was silently mutated to UUUCCTG in a similar way using site-directed mutagenesis and HPLC purified primers (table S3) to produce only 6K CHIKV - ΔTF CHIKV construct.

### CHIKV infection

For all CHIKV infections, HEK293T cells were seeded 24 h prior to infection by CHIKV (strain 119067) (70) at a MOI of 0.1. CHIKV was added to the cells post phosphate-buffered saline (PBS) washing for adsorption. After 90 min, cells were again washed with PBS, and complete media was added. Post-infection, cells were incubated for 24-48 hours under normal cell growth conditions.

### Mutated CHIKV generation by reverse genetics

A mutant Chikungunya virus (CHIKV) infectious clone lacking translational frameshifting (ΔTF-CHIKV) was generated using a previously described mCherry-CHIKV infectious cDNA clone as the parental backbone (71). Site-directed mutagenesis (SDM) was performed to disrupt the ribosomal frameshift site responsible for translational frameshifting, without altering the encoded open reading frame or the resulting amino acid sequence. Mutagenic primers (table S3) were designed accordingly and synthesized with HPLC purification. PCR-based SDM was performed using KOD Hot Start DNA Polymerase (Merck) according to the manufacturer’s instructions. The amplified products were processed according to standard cloning procedures to obtain the mutated plasmid. All introduced mutations were confirmed by Sanger sequencing, ensuring the integrity of the viral genome and the absence of unintended secondary mutations Vero E6 cells were seeded in a 6-well plate to reach 60% confluency. Cells were then transfected with 5µg of CHIKV plasmid (WT CHIKV) and the mutant generated (ΔTF CHIKV) with polyethyleneimine (PEI) in a ratio of 1:5. The cell supernatant was collected 48 hours post-transfection, centrifuged at 2000 rpm for 10 minutes at 4°C. The supernatant was carefully removed, aliquoted, and stored at -80°C. For virus propagation, Vero E6 cells were cultured up to 80% confluency in DMEM supplemented with 10% FBS. Before infection, cells were incubated in incomplete media for 2-3 hours. Thereafter, cells were infected with 0.01 MOI of CHIKV and observed for maximum cytopathic effect (40-48 hours post-infection). The supernatant was collected and centrifuged at 2,000g for 10 minutes at 4°C to remove cell debris. The virus supernatant was carefully separated without disturbing the cell pellet and stored at -80°C.

### Virus propagation and purification

Vero cells were cultured in DMEM (Gibco), supplemented with 10% fetal bovine serum (Gibco) and 1% Penicillin-Streptomycin (Gibco) at 37 °C in 5 % CO2 environment. Cells were subsequently infected with ΔTF CHIKV in serum-free DMEM medium. The culture supernatant was collected 48 hours after infection. ΔTF CHIKV particles were precipitated using 10% PEG 8000. The pellet was resuspended in 20 mM HEPES (pH 7.5), 150 mM NaCl, and 1 mM EDTA. The resuspended pellet was layered onto a 30-60% sucrose gradient and centrifuged at 35,000 rpm in a SW41Ti rotor for 2 hours at 4 °C. The viral band was subsequently pelleted on a 30% sucrose cushion at 40,500 rpm in a SW41Ti rotor for 1 hour at 4 °C. The pellet was resuspended in 20 mM HEPES (pH 7.5). 150 mM NaCl, 1 mM EDTA. The virus concentration was assessed using BCA.

### Western blotting

HEK293T cells infected with CHIKV were washed with PBS and lysed with RIPA lysis buffer supplemented with a protease inhibitor cocktail (ThermoFisher Scientific) as per the manufacturer’s protocol. Total protein was quantified using the bicinchoninic acid (BCA) assay, followed by SDS-PAGE and western blotting with a primary antibody against Scribble (1:500, Santa Cruz sc55543), β -actin (1:1000, Cell signaling Technology 4967), β -tubulin (1:200, Santa Cruz sc55529), CHIKV E2 (1:3000, The Native antigen company MAB12130-200), and EGFP (1:3000, Bio Bharati BB-AB0065). Densitometric analysis was performed to quantify the protein levels in western blots using the Fiji software as described elsewhere (72,73).

### TF peptide production and generation of specific antibodies

A 15-amino acid peptide corresponding to the C-terminal region of TF was chemically synthesized and coupled to a carrier protein, Keyhole Limpet Hemocyanin (KLH), via the N-terminus (ThermoFisher Scientific). The KLH-TF peptide was dissolved in DMSO and stored at –20°C until use.

Polyclonal sera against TF were raised by immunizing two New Zealand white rabbits, aged 3-4 months, with the KLH-TF peptide emulsified in complete/incomplete Freund’s adjuvant (CFA/IFA) as per the following protocol. On Day 0 (priming), 1 mg of KLH-TF peptide was emulsified with CFA and injected subcutaneously into the rabbits. Subsequently, four boosts (on Days 21, 42, 63, 84) were given with 300 µg of the peptide emulsified in IFA intramuscularly. Sera were collected two weeks post-immunization from boost 2 onwards and checked for anti-TF antibodies using immunoblotting.

### Immunoprecipitation

HEK293T cells were infected with CHIKV at an MOI of 0.1. Cell extracts were prepared 24 hours post-infection. A polyclonal antibody against Scribble (Invitrogen PA5-54821) was added overnight to precleared cell extracts at 4 °C on a rocker, followed by incubation with protein A/G agarose beads (Santa Cruz sc-2003) for 1 hour at 4 °C with gentle mixing. The beads were washed thrice with lysis buffer (Tris HCl – 20 mM, KCl – 120 mM, EDTA – 0.2 mM, DTT – 1 mM, and protease inhibitor cocktail). Post-washing, the beads were resuspended in SDS-PAGE buffer, and the samples were analyzed in immunoblots with anti-Scribble and anti-TF antibodies.

### MBP-tagged protein production in bacteria

Protein production was carried out as previously described (19). Briefly the cDNA corresponding to the TF and 6K regions of CHIKV were cloned into the pMALc6T vector (New England Biolabs), in conjunction with N-terminal Histidine and Maltose binding protein (MBP) tags, and a TEV protease cleavage site. Recombinant expression of MBP-tagged TF and 6K proteins was induced by the addition of Isopropyl ß-D-1-thiogalactopyranoside in *E. coli* Rosetta pLysS cells at 16 °C at an OD 600 of 0.6. The expressed proteins were 2-step purified by Ni-NTA affinity chromatography followed by size exclusion chromatography (SEC) in a buffer containing 25 mM HEPES, 100 mM NaCl, and the detergent n-Dodecyl β-D maltoside (DDM).

### Confocal microscopy

HEK293T cells were cultured on glass coverslips and infected with CHIKV at an MOI of 0.1. Twenty-four hours post-infection, cells were washed with chilled PBS and fixed with 4% paraformaldehyde (PFA) for 10 min, followed by permeabilization with 0.1 % Triton X-100 in PBS for 5 min and blocked with 5% bovine serum albumin for 60 min. Immunostaining was performed at 4 °C for Scribble, Ubiquitin, and CHIKV proteins using anti-Scribble (1:500, Invitrogen PA5-54821), anti-Ubiquitin (1:300, Santa Cruz sc-8017), anti-E2 (1:500, The Native Antigen Company MAB12130-200), and anti-capsid (1:200, The Native Antigen Company MAB12131-200) antibodies. Post-primary antibody incubation, Alexa fluor-labelled appropriate secondary antibodies were added for 60 min. All washes were carried out using chilled PBS. Nuclei were labelled with DAPI (Invitrogen D1306) for 10 min, and cells were mounted using Prolong Glass Antifade Mountant (Invitrogen P36982). The slides were dried for a few minutes and sealed with nail paint. Imaging was performed on a confocal microscope (Leica TCS SP8) at a magnification of 63X using the oil immersion objective.

For transfection of pEGFPC1 constructs, HEK293T cells were seeded and grown at 37°C for 24 h, followed by the addition of plasmid DNA in conjunction with lipofectamine2000 (Invitrogen) according to the manufacturer’s protocol. Briefly, DNA and lipofectamine (1:3 w/v) were diluted in Opti-MEM (Gibco) and incubated at room temperature (RT) for 20 min. The DNA-lipofectamine complexes were used to replace the cellular growth medium for 5 h. Twenty-four hours post-transfection, cells were washed thrice with PBS, fixed with 4% PFA, and processed as above. Co-localization was estimated as the percentage of qualified double-positive cells relative to the total number of infected or transfected cells as visualized in four distinct fields.

### Confocal data analysis

Line intensity profiles were generated from confocal images by drawing a line across regions of interest, and fluorescence intensity values for each channel were plotted along the length of the line. Image analysis was performed in Fiji (72). Prior to analysis, images were background-subtracted using a rolling-ball algorithm (radius 50 pixels). Scribble puncta were identified by thresholding the Scribble channel and used to generate region-of-interest masks. Colocalization was quantified using the Coloc2 plugin by calculating Manders’ overlap coefficients with Costes’ automatic thresholding. Colocalization analysis was performed only in conditions displaying discrete Scribble puncta. For each condition, at least 10 cells from three independent experiments were analyzed (74). Data are presented as mean ± SEM.

### Mass spectroscopy

#### Sample Preparation

25μg protein sample was first reduced with 5 mM TCEP and subsequently alkylated with 50 mM iodoacetamide. The protein was then digested with Trypsin at a 1:50 Trypsin-to-lysate ratio for 16 hours at 37°C. After digestion, the mixture was purified using a C18 silica cartridge and then concentrated by drying in a speed vac. The resulting dried pellet was resuspended in buffer A, which consists of 2% acetonitrile and 0.1% formic acid.

#### Mass Spectrometric Analysis of Peptide Mixtures

For mass spectrometric analysis, all the experiments were performed on an Easy-nLC-1000 system (Thermo Fisher Scientific) coupled with an Orbitrap Exploris 240 mass spectrometer (Thermo Fisher Scientific) and equipped with a nano electrospray ion source. 1ug of phosphopeptides sample was dissolved in buffer A containing 2% acetonitrile/0.1% formic acid and resolved using Picofrit column (1.8-micron resin, 15cm length). Gradient elution was performed with a 0–38% gradient of buffer B (80% acetonitrile, 0.1% formic acid) at a flow rate of 500nl/min for 96mins, followed by 90% of buffer B for 11 min and finally column equilibration for 3 minutes. Orbitrap Exploris 240 was used to acquire MS spectra under the following conditions: Max IT = 60ms, AGC target = 300%; RF Lens = 70%; R = 60K, mass range = 375−1500. MS2 data was collected using the following conditions: Max IT= 60ms, R= 15K, AGC target 100%. MS/MS data was acquired using a data-dependent top20 method dynamically choosing the most abundant precursor ions from the survey scan, wherein dynamic exclusion was employed for 30s.

#### Data Processing

Sample was processed and RAW files generated was analyzed with Proteome Discoverer (v2.5) against UniProt reference database. For dual Sequest and Amanda search, the precursor and fragment mass tolerances were set at 10 ppm and 0.02 Da, respectively. The protease used to generate peptides, i.e. enzyme specificity was set for trypsin/P (cleavage at the C terminus of “K/R: unless followed by “P”). Carbamidomethyl on cysteine as fixed modification and oxidation of methionine and N-terminal acetylation were considered as variable modifications for database search. Both peptide spectrum match and protein false discovery rate were set to 0.01 FDR.

### shRNA mediated knockdown of Scribble

HEK293T cells were seeded and grown at 37 °C for 24 h, and then the Scribble short hairpin RNA (shRNA) containing plasmid DNAs were transfected with lipofectamine 2000 as described earlier. Four different shRNAs were used in different conditions, either alone or in combination, to optimize Scribble knockdown. The effect of the knockdown was analyzed 36 hours post-transfection using western blot analysis and the results were quantified via densitometry. Two conditions, shRNA.C1: AAACCACAAAATAGAGTCT, and shRNA.C2: a combination of AAACCACAAAATAGAGTCT and TGTACAAACATCACTAGTT, produced maximum and consistent knockdown of ∼50 %, and were utilized for further studies.

### MTT assay

Assays with the tetrazolium dye 3-[4, 5-dimethylthiazol-2-yl]-2, 5-diphenyl tetrazolium bromide (MTT) were performed as described in the product’s manual (Himedia) with slight modifications. Briefly, 10 µl MTT (5 mg/ml, Himedia) was added to cells, and the formazan crystals were dissolved in dimethyl sulfoxide after 30 min. The absorbance measurements were recorded at 570 and 690 nm.

### Quantitative real-time polymerase chain reaction (qRT-PCR)

CHIKV-infected HEK293T cells were washed with PBS, harvested in Trizol (Takara), and stored at -80 °C. RNA was isolated from the samples as per the manufacturer’s protocol, quantitated using NanoDrop (Thermo Scientific), and 5 µg RNA was reverse transcribed into cDNA using a cDNA synthesis kit (Takara Bio) as per the manufacturer’s instructions. The cDNA was used to analyze human Scribble and CHIKV E1 expression levels by qRT-PCR using gene-specific primers (Supplementary Table S3). PCR reactions were conducted on an ABI 7500 FAST mastercycler, and the relative quantification of gene expression was carried out using the ΔΔCt method with GAPDH as the housekeeping gene.

### Plaque assay

Vero cells were seeded in a six-well plate in DMEM with 10% FBS (Gibco) and antibiotics (Corning) and incubated for 24 h at 37°C, 5% CO_2_. The cell growth media was then replaced with serum-free media, and cells were incubated for 4-6 h. Subsequently, 10-fold dilutions (10^-5^ to 10^-7^) of the virus-containing culture supernatants of CHIKV-infected HEK293T cells, in serum-free media, were used to replace the cellular growth media. Virus particles in the diluted culture supernatant were allowed to infect Vero cells for 1 h with shaking every 15 min. Post 1 h, the cells were washed with PBS, and 3 ml of an overlay mix of 2% low melting agarose in 2X DMEM was added to each well. The agarose was allowed to solidify at 4°C for 15 min, and cells were then incubated at 37°C for 48 h. For plaque visualization, cells were fixed with 10% formaldehyde for 4-5 h at room temperature and stained with crystal violet for 5 min, followed by washing with tap water. The plate was allowed to dry for a few minutes, and the plaques were counted. The plaque-forming units (PFU) were calculated using the following formula: PFU/ml = (number of plaques x dilution factor)/(inoculum volume).

### Cryo-electron microscopy and 3D image reconstruction

ΔTF CHIKV particles were prepared for cryo-electron microscopy by applying 3.5 µl of sample onto freshly glow-discharged 300-mesh Quantifoil Au R 2/2 grids. The grids were incubated for 30 s, blotted for 3 s at 100% humidity, and plunge-frozen in liquid nitrogen–cooled liquid ethane using a Vitrobot (Thermo Fisher Scientific). Vitrified grids were imaged on a 300 keV Thermo Fisher Titan Krios equipped with a BioQuantum energy filter and K3 direct electron detector (Gatan). Movies were collected at a nominal magnification of 81,000×, corresponding to a pixel size of 1.1 Å/pixel.

Movie frames were imported into cryoSPARC v4.4.0 (75) for data processing. Patch-based motion correction and CTF estimation were performed, followed by automated particle picking using blob picking. A total of 135,422 particles were extracted with a box size of 850 pixels and subjected to multiple rounds of 2D classification, *ab initio* model generation, and 3D classification to remove damaged and poorly resolved particles. The final dataset of 84,104 particles was refined with imposed icosahedral (I) symmetry. Density maps were visualized using UCSF Chimera (76). A side-by-side reconstruction of WT CHIKV was carried out using a similar workflow.

## Acknowledgements

The work was funded by the Science and Engineering Research Board, Government of India (grant number SPF/2021/000115) and Indian Council of Medical Research (IIRPIG-2024-01-00529). The authors thank the Sophisticated Analytical & Technical Help Institutes (SATHI) foundation and Central Research Facility (CRF), IIT Delhi for the National CryoEM facility and confocal microscopy facility respectively; and the High-Performance Computational Facility (HPC), IIT Delhi for simulation studies. The mcherry CHIKV infectious clone was a kind gift from Prof. Andres Merits from the University of Tartu, Estonia to SB. The authors extend their thanks to Abhishek Trivedi (SATHI IIT Delhi) and Dr. Anjali Kapoor (CRF IIT Delhi) for their assistance during cryoelectron microscopy data collection and confocal microscopy imaging respectively. The authors also thank Prof. Manoj Menon, KSBS, IIT Delhi for providing antibodies. RK and PT would also like to acknowledge IIT Delhi for providing fellowship support.

## Author contributions

Conceptualization: RK, DD, PT, MB; Methodology: PT, RK, DD, MB; Investigation: RK, PT, DD, YR, SBK, B, SA, SYM, SJ, KS; Resources: SB, MB; Writing-original draft: RK, PT, DD, MB; Writing- review & editing: RK, PT, MB; Visualization: RK, PT, MB; Supervision: SB, MB. Project administration: MB; Funding acquisition: MB.

## Conflict of interest

The authors declare no conflict of interest.

## Data availability

Cryo-EM density maps of ΔTF CHIKV and WT CHIKV were deposited in the Electron Microscopy Data Bank under the accession codes EMD-69164 and EMD-69185, respectively.

## Notes

### Competing Interest Statement

The authors have declared no competing interest.

### Summary of Updates

This version of the manuscript has been revised to include additional experimental results that further strengthen our findings on the role of the alphavirus TF protein in viral propagation and modulation of host cell boundaries. The major update in this version is the inclusion of structural analysis of the Chikungunya virus. We have now added cryo-electron microscopy data describing the structure of both the wild-type virus and the TF mutant virus. These structural data provide new insights into the impact of TF on viral architecture and particle organization.

