## Supplementary information for "The alphavirus TF protein plays a critical role in promoting viral propagation by altering cell-cell boundaries"

**Supplementary figures**

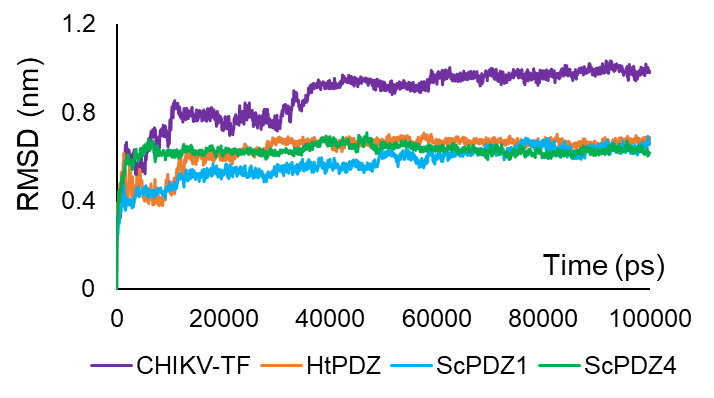

**Fig. S1. RMSD plot for TF interactions with Scribble and HTRA1**

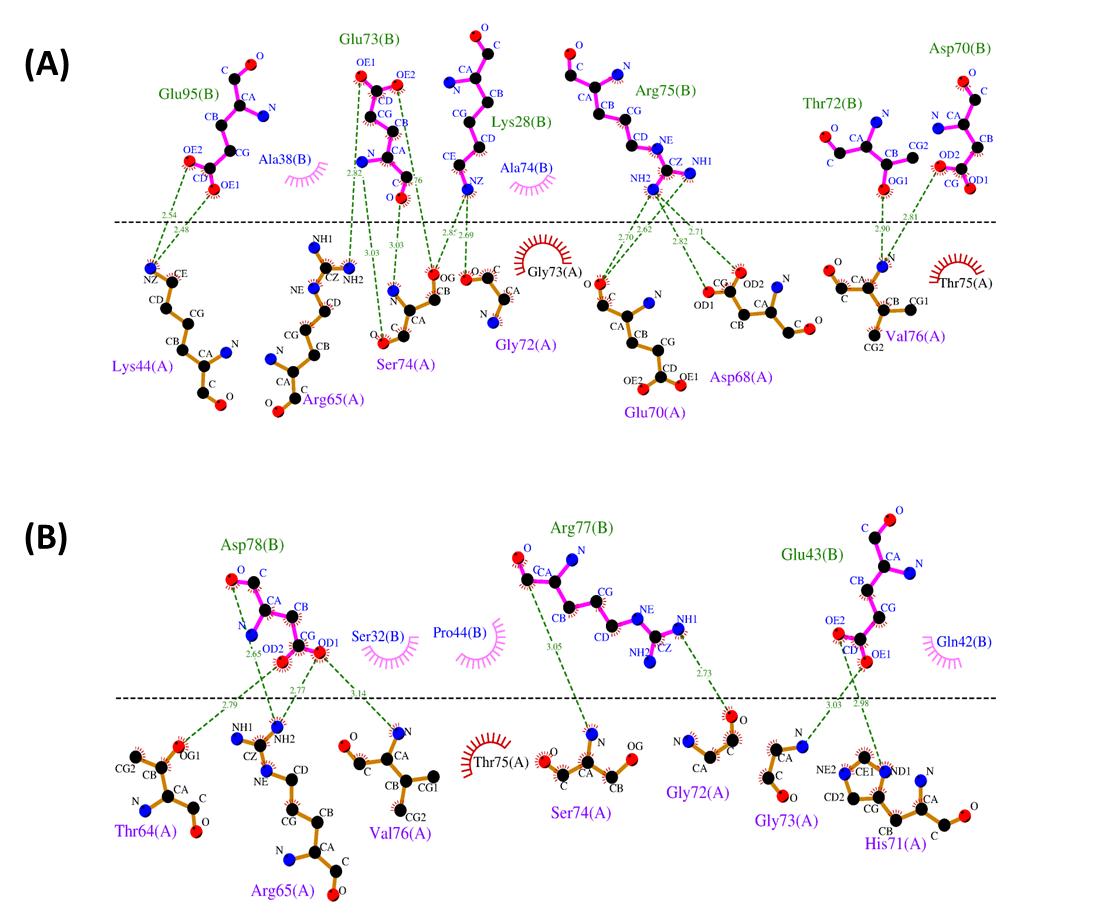

**(A)**

**(B)**

**Fig. S2. Predicted interactions between TF and Scribble PDZ2 (A), and PDZ3 (B).**

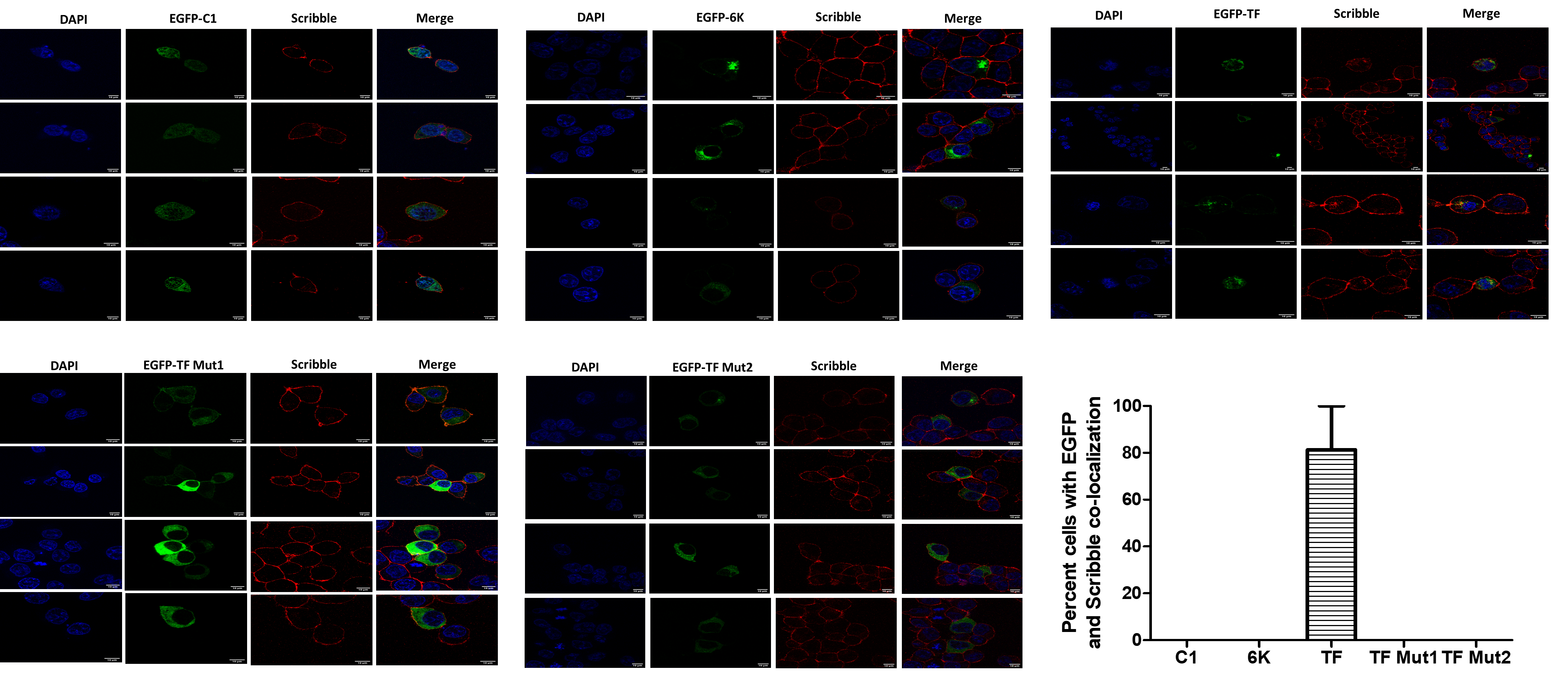

**Fig. S3. Quantification of co-localization of EGFP-tagged proteins and Scribble in transfected HEK293T cells.** HEK293T cells were transfected with pEGFPC1 vector or pEGFPC1 plasmids carrying TF or 6K. Twenty-four hours post-transfection, cells were fixed, and immunostaining was performed for Scribble (red). For each construct, 4 fields were analyzed. Data represents mean ± SEM for cells visualized in 4 different fields. Scale - 10μm

Manders’ coefficient

| C1-TF expressing cells | Manders |
| --- | --- |
| 1 | 0.871 |
| 2 | 0.712 |
| 3 | 0.78 |
| 4 | 0.826 |
| 5 | 0.69 |
| 6 | 0.71 |
| 7 | 0.72 |
| 8 | 0.874 |
| 9 | 0.59 |
| 10 | 0.721 |

| C1-TF expressing cells | Mean |
| --- | --- |
| 1.01723 | 0.7494 |

**Fig. S4. Quantification of colocalization between GFP-TF and Scribble using Manders’ coefficient.** Colocalization between GFP-TF and Scribble was quantified by calculating Manders’ overlap coefficients for individual cells. Each dot represents one cell. The orange dot indicates the mean Manders’ coefficient, and error bars represent SEM. Analysis was performed on Scribble puncta identified by thresholding, using the Coloc2 plugin in Fiji with Costes automatic thresholding.

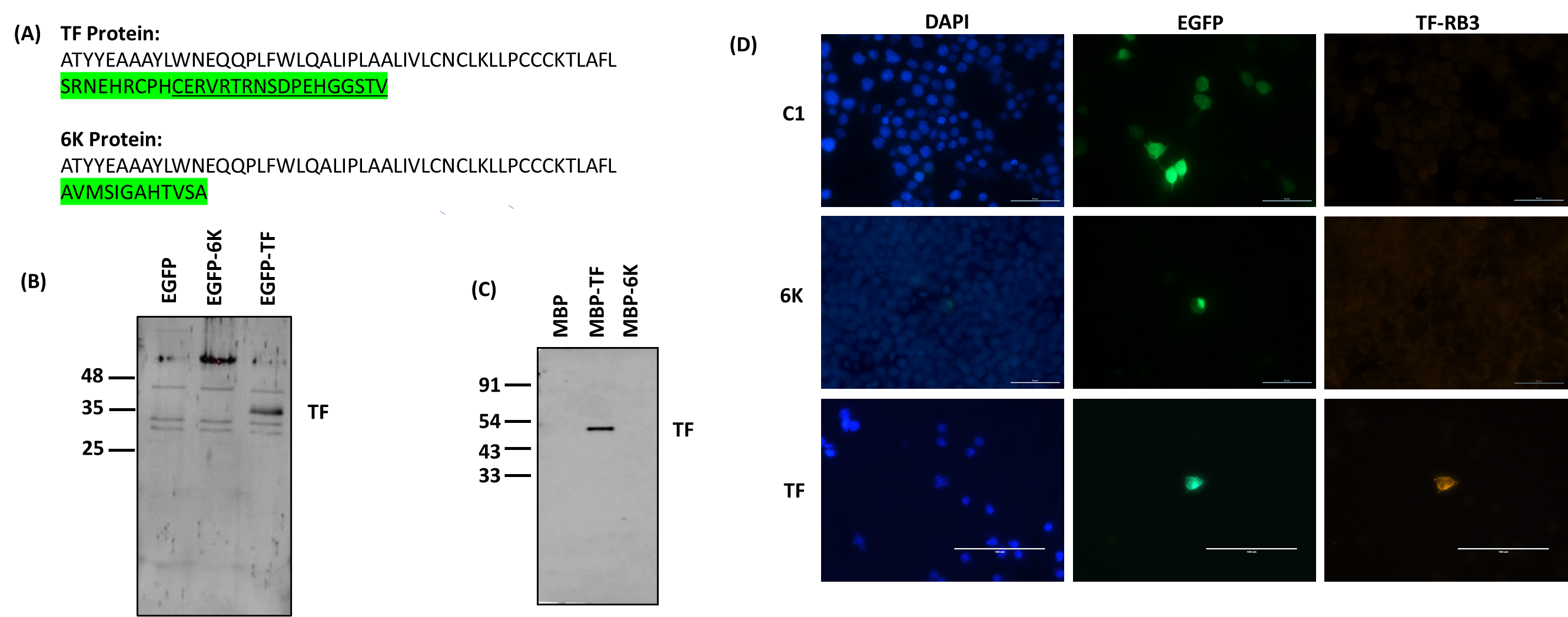

**Fig. S5. Assessment of TF antisera.** TF peptide sequence used for generating anti-TF antibodies is underlined. The region highlighted in green represents the non-identical region between TF and 6K (A). Western blot of HEK293T cells transfected with pEGFPC1 vector or pEGFPC1 plasmids carrying TF or 6K (B). Western blot of MBP, MBP-TF, and MBP-6K purified protein with the anti-TF antisera (C). Immunostaining of TF in pEGFPC1 constructs transfected HEK293T cells (D).

Manders’ coefficient

| TF expressing cells |  | Mean |
| --- | --- | --- |
| 1 | 0.72 | 0.77 |
| 2 | 0.89 |  |
| 3 | 0.59 |  |
| 4 | 0.82 |  |
| 5 | 0.68 |  |
| 6 | 0.56 |  |
| 7 | 0.91 |  |
| 8 | 0.71 |  |
| 9 | 0.88 |  |
| 10 | 0.94 |  |

**Fig. S6. Quantification of colocalization between TF and Scribble using Manders’ coefficient.** Colocalization between TF and Scribble in CHIKV-infected cells was quantified by calculating Manders’ overlap coefficients for individual cells. Each dot represents one cell. The orange dot indicates the mean Manders’ coefficient, and error bars represent SEM. Analysis was performed on Scribble puncta identified by thresholding, using the Coloc2 plugin in Fiji with Costes automatic thresholding.

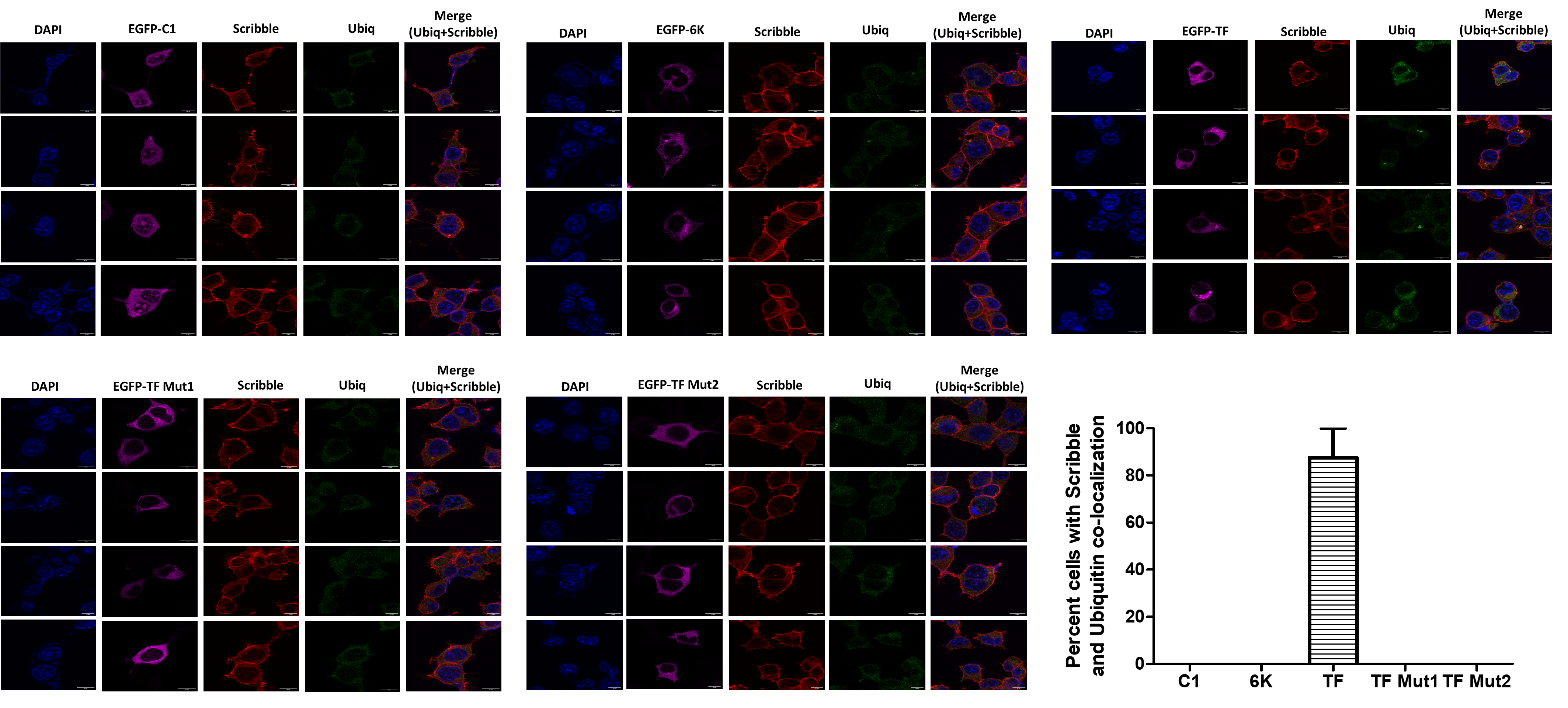

**Fig. S7. Quantification of co-localization of Scribble and Ubiquitin in transfected HEK293T cells.** Effect of TF on Scribble ubiquitination. HEK293T cells were transfected with pEGFPC1 vector or pEGFPC1 plasmids carrying 6K, TF and TF mutants. Twenty-four hours post-transfection, cells were fixed, and immunostaining was performed for Scribble (red) and Ubiquitin (green). For each construct, 4 fields were analyzed. Data represents mean ± SEM for cells visualized in 4 different fields. Ubiq: Ubiquitin. Scale - 10μm

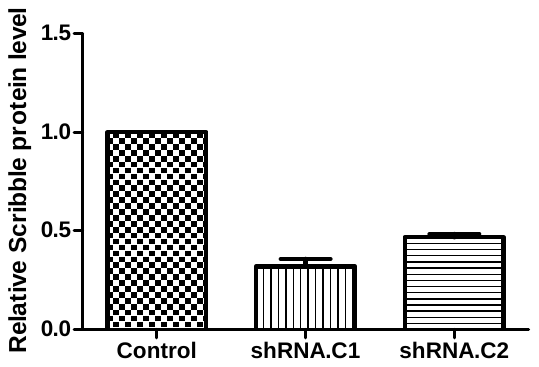

**(B)**

**(A)**

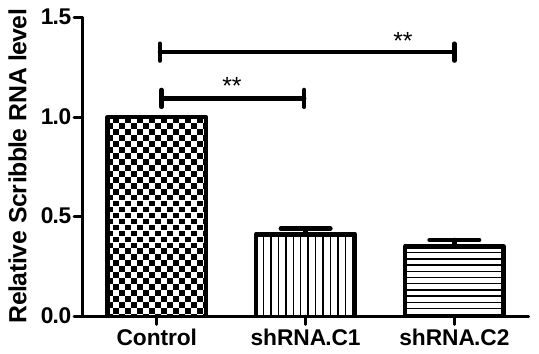

**(C)**

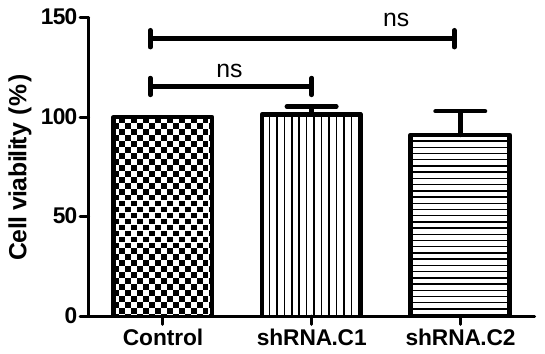

**Fig. S8. Effect of Scribble knockdown on CHIKV infection.** Scrib shRNAs were transfected into HEK293T cells, and knockdown was confirmed 36 hours post-transfection using western blot and quantitated using densitometry (A). Scribble knockdown was also confirmed at RNA level using real-time PCR (B). The effect of Scribble knockdown on cell viability was assessed using MTT assay (C).

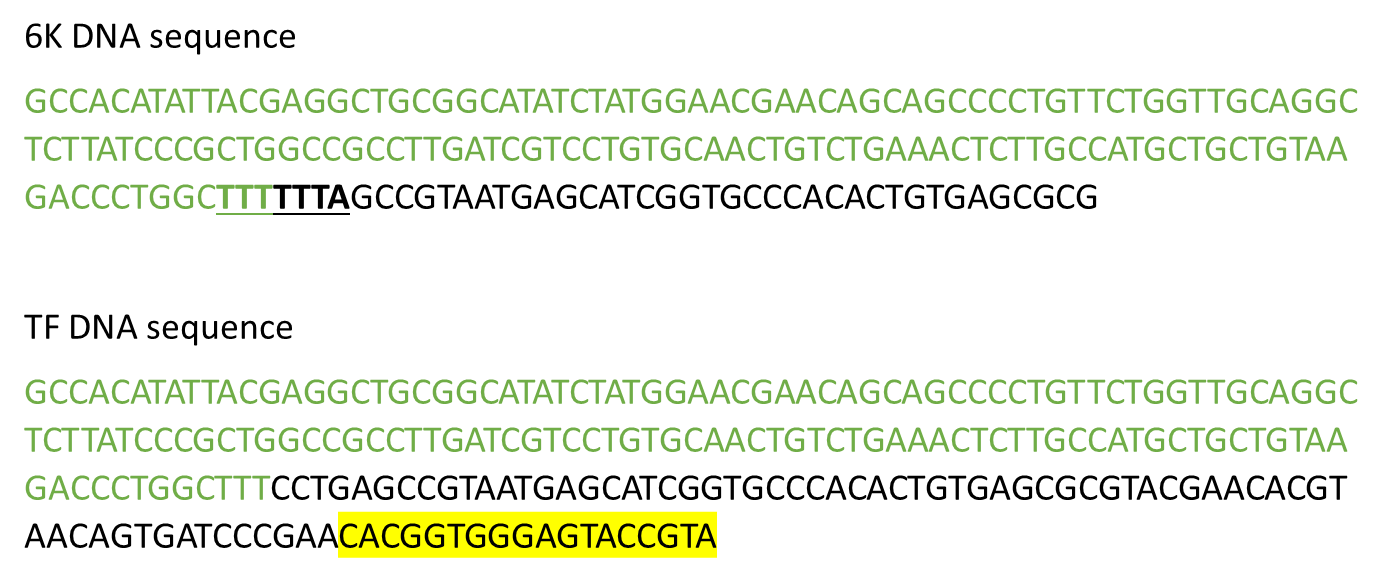

**Fig. S9. CHIKV 6K and TF cDNA sequences.** The sequences that are common to both 6K and TF are written in green. The slippery site in 6K is bold and underlined. The predicted PDZ binding motif in TF is highlighted in yellow.

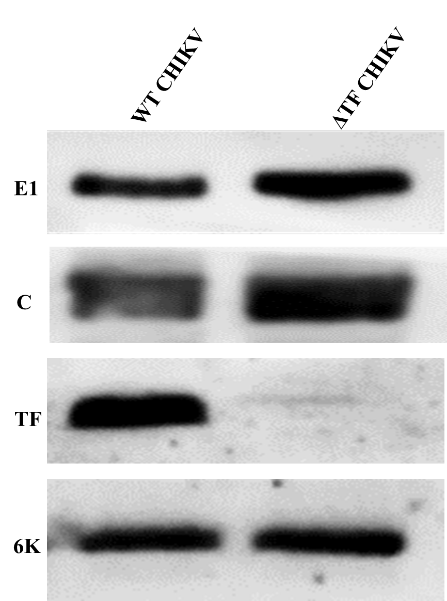

**Fig. S10. Glycoprotein expression.** Western blot of HEK293T cell lysate infected at 0.1 MOI with WT CHIKV and **Δ**TF CHIKV showing the expression of E1, Capsid, TF and 6K protein.

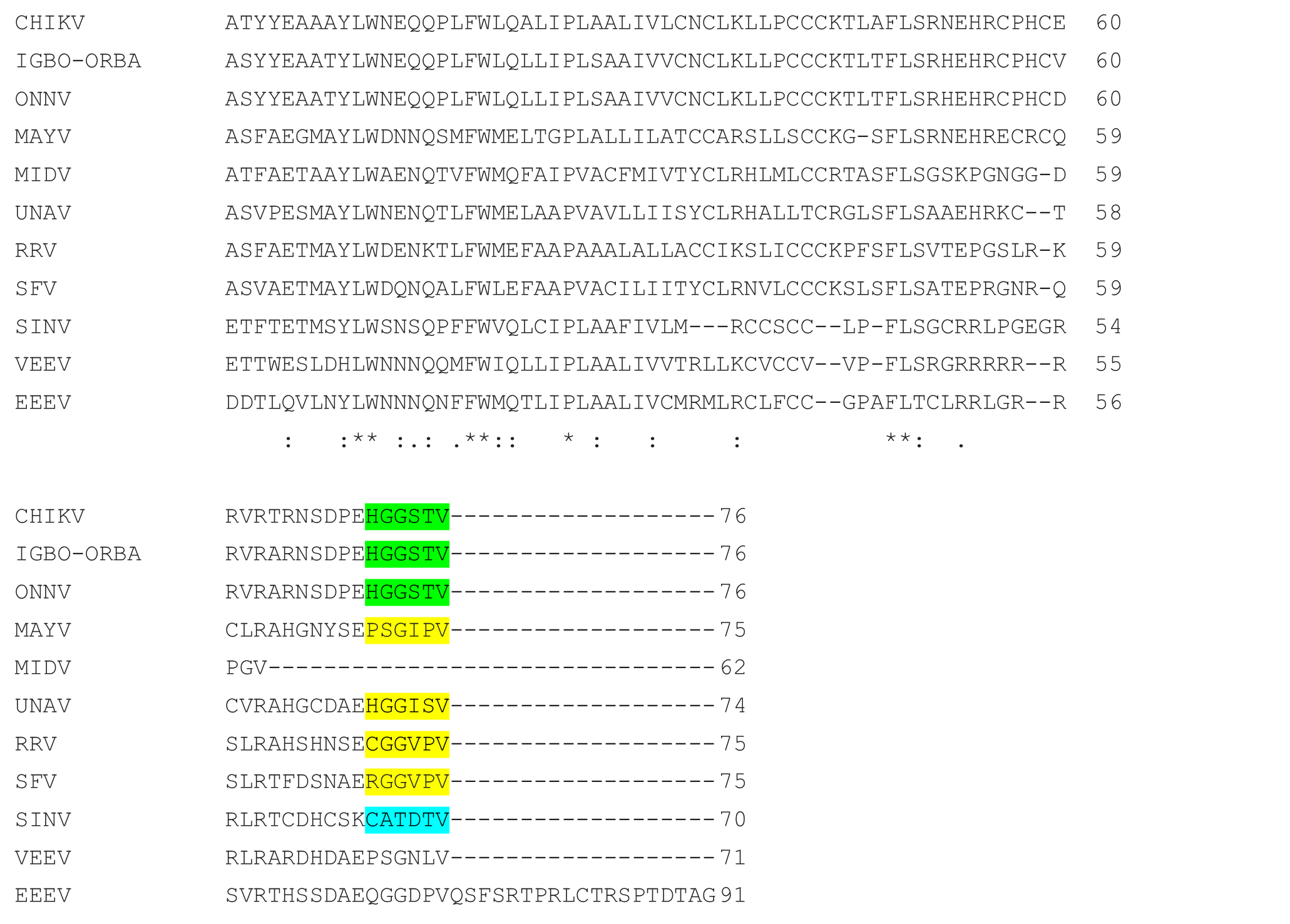

**Fig. S11. PDZ binding motif in TF sequences from alphaviruses.** CLUSTAL O (1.2.4) multiple sequence alignment of CHIKV (strain 37997) TF sequence with other alphaviruses. The TF sequence for other alphaviruses were taken from Firth et al., 2008. The regions highlighted in green, yellow, and blue represent class I, II, and III PBM, respectively, as predicted by the ELM software [45]. ‘*’ represents identical residues in the alignment. ‘:’ represents strongly similar residues. ‘.’ represents weakly similar residues.

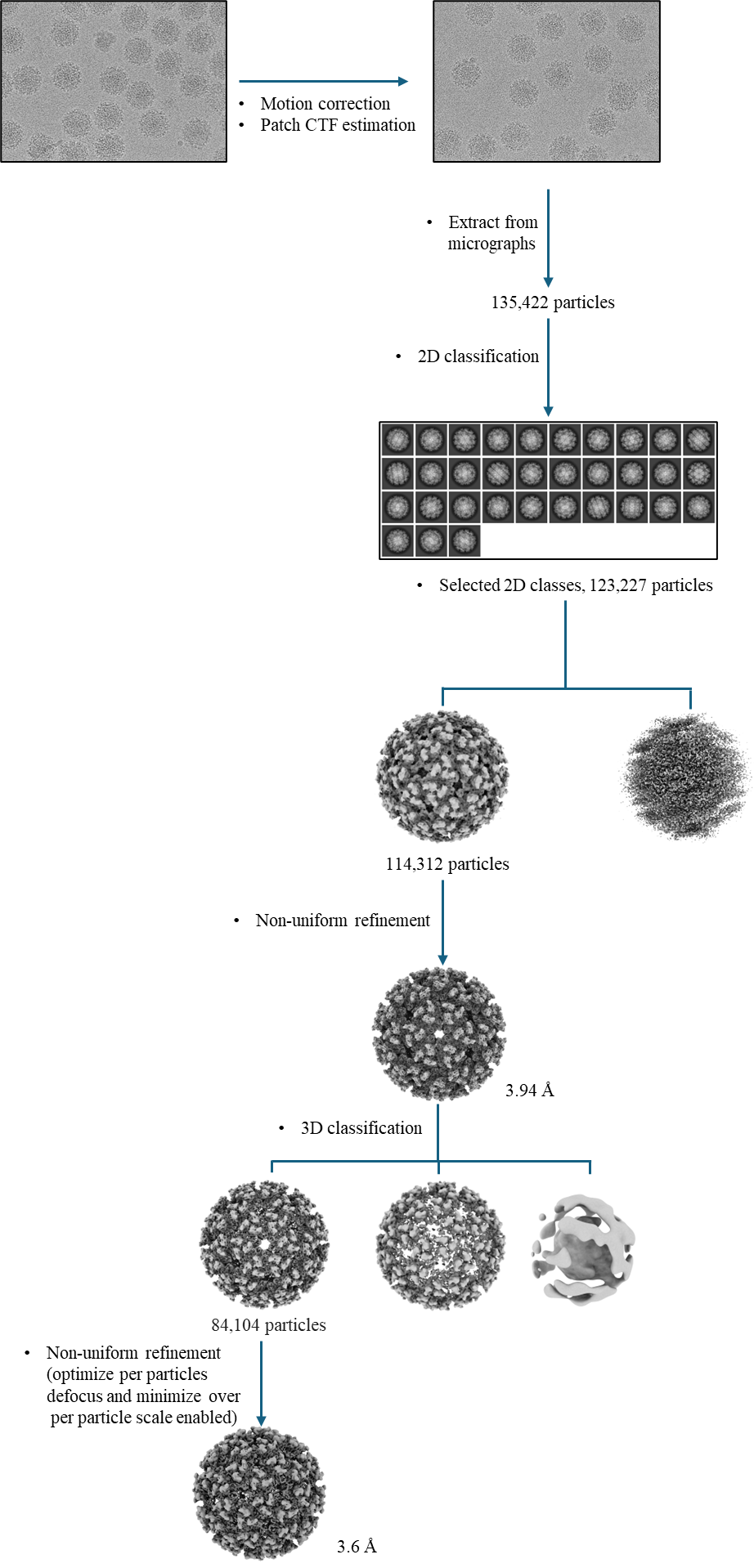

**Figure S12: Data processing workflow in cryoSPARCSupplementary tables**

**Table S1. Binding energies for TF interactions**

| **System** | **Binding Energy (kJ/mol)** | **Van der waals Energy (kJ/mol)** | **Electrostatic Energy (kJ/mol)** | **Polar Solvation Energy (kJ/mol)** | **SASA Energy (kJ/mol)** |
| --- | --- | --- | --- | --- | --- |
| TF-ScPDZ1 | -530.075 ± 71.564 | -652.267 ± 32.621 | -886.300 ± 81.887 | 1065.847 ± 98.420 | -57.354 ± 3.442 |
| TF-ScPDZ2 | -208.128 ± 5.381 | -359.047 ± 1.804 | -797.578 ± 5.857 | 986.052 ± 7.766 | -37.597 ± 0.237 |
| TF-ScPDZ3 | -258.072 ± 7.180 | -391.921 ± 2.572 | -969.140 ± 8.420 | 1145.149 ± 12.746 | -42.175 ± 0.342 |
| TF-ScPDZ4 | -349.404 ± 73.230 | -671.951 ± 42.247 | -617.994 ± 91.476 | 1002.387 ± 121.825 | -61.846 ± 4.173 |
| TF-HtPDZ | -91.844 ± 75.366 | -562.245 ± 30.714 | -417.963 ± 61.048 | 940.652 ± 97.128 | -52.288 ± 3.966 |

**Table S2. Prediction of PBM in alphavirus TF using the ELM software** (25)

| **Alphavirus** | **Region** | **PBM** | **Class** |
| --- | --- | --- | --- |
| SINV | 65-70 | CATDTV | Class_3 |
| ONNV | 71-76 | HGGSTV | Class_1 |
| UNAV | 69-74 | HGGISV | Class_2 |
| RRV | 70-75 | CGGVPV | Class_2 |
| IGBO-ORBA | 71-76 | HGGSTV | Class_1 |
| MAYV | 70-75 | PSGIPV | Class_2 |
| SFV | 70-75 | RGGVPV | Class_2 |
| EEEV | NA |  |  |
| VEEV | NA |  |  |
| MIDV | NA |  |  |

**Table S3: List of primers**

| **Purpose** | **Primer** | **Sequence (5’ → 3’)** |
| --- | --- | --- |
| DNA sequencing | EGFP-CF | AGCACCCAGTCCGCCCTGAGC |
| RT-PCR | hScribble_Forward | CATCGGGGACTGTGAGAACC |
| RT-PCR | hScribble_Reverse | TCCACGTTGAGGTTGGTCAG |
| RT-PCR | CHIKV_E1_Forward | AAGTACACTGTGCAGCTGAGT |
| RT-PCR | CHIKV_E1_Reverse | GCATAGCACCACGATTAGAATC |
| SDM | 6K Silent_Forward | TGTAAGACCCTGGCTTTCCTGGCCGTAATGAGCATC |
| SDM | 6K_Silent_Reverse | GATGCTCATTACGGCCAGGAAAGCCAGGGTCTTACA |
| SDM | TF Mut1_Forward | CCCGAACACGGTGCCGCTACCGTATAAAAGC |
| SDM | TF Mut1_Reverse | GCTTTTATACGGTAGCGGCACCGTGTTCGGG |
| SDM | TF Mut2_Forward | CAGTGATCCCGAAGCCGCTGCCGCTACCGTATA |
| SDM | TF Mut2_Reverse | TATACGGTAGCGGCAGCGGCTTCGGGATCACTG |
| SDM | ΔTF_Forward | TGTAAAACGTTGGCTTTCCTGGCCGTAATGAGCGTC |
| SDM | ΔTF_Reverse | GACGCTCATTACGGCCAGGAAAGCCAACGTTTTACA |
